# Global Stability and Tipping Point Prediction in a Coral-Algae Model Using Landscape-Flux Theory

**DOI:** 10.1101/2024.12.17.627631

**Authors:** Li Xu, Denis Patterson, Simon Levin, Jin Wang

**Author notes:** **Correspondence:** Simon Levin and Jin Wang.

## Abstract

Coral reef ecosystems are remarkable for their biodiversity and ecological significance, exhibiting the capacity to exist in different stable configurations with possible abrupt shifts between these alternative stable states. This study applies landscape-flux theory to analyze how these complex systems behave when subjected to random environmental disturbances. We use this theory to formulate and investigate several early warning indicators of ecosystem transitions in a well-known coral-reef model. We studied a number of specific indicators, including the average flux (the driving force when the system is out of equilibrium), the entropy production rate, the nonequilibrium free energy and the time irreversibility of the cross-correlation functions. These indicators demonstrate a distinctive advantage when compared to classical indicators based on the phenomenon of critical slowing down; they exhibit turning points midway between two bifurcations, enabling them to forecast transitions in both directions substantially earlier than conventional methods. In contrast, early warning indicators based on the critical slowing down phenomenon typically only become apparent when the system approaches the actual bifurcation or tipping point(s). Our findings offer improved tools for anticipating critical transitions in coral reef and other at-risk ecosystems, with the potential to enhance conservation and management strategies.

## 1 Introduction

These complex systems provide critical ecological functions and substantial economic value through coastal protection and support of fish and marine biodiversity(Mcmanus and Polsenberg, 2004; Hughes et al., 2007; Mccook et al., 2001; Dudgeon et al., 2010). However, coral reefs globally are confronting multiple challenges and experiencing serious threats to their abundance, diversity, structural integrity, and ecological functioning(Mumby et al., 2007; Li et al., 2014; Mcmanus et al., 2018; Nes et al., 2016). The degradation of coral reef ecosystems results from a synergistic combination of anthropogenic pressures (including overfishing and pollution) and natural disturbances (such as disease outbreaks, hurricanes, and coral bleaching events).

The magnitude of this decline is striking-average hard coral cover in the Caribbean Basin has plummeted from approximately 50% to merely 10% over just three decades since 1977(Diko, 2010; Pandolfi et al., 2003). While algal proliferation rarely causes direct coral mortality, these organisms compete with corals for essential resources such as space and light, contributing to the death of established coral colonies. Furthermore, algae impede coral recruitment and regeneration, thereby undermining the capacity of coral populations to recover from environmental stressors(Mumby et al., 2007; Li et al., 2014; Mcmanus et al., 2018; Nes et al., 2016). The most dramatic illustration of such transformation is observed in Caribbean reefs, which have undergone a profound shift to an alternative stable state dominated by algal cover(Mumby et al., 2007; Li et al., 2014; Mcmanus et al., 2018; Nes et al., 2016). This striking ecological transition represents one of the most well-documented examples of regime shifts in marine ecosystems, fundamentally altering both reef structure and function.

Human land use activities have increased oceanic nutrient loading, promoting excessive algal growth in marine ecosystems. Historically, herbivorous fish have played a crucial role in regulating algal biomass(Mumby et al., 2007; Li et al., 2014). However, widespread overfishing has significantly reduced populations of important herbivores such as parrotfish. These herbivores primarily consume algae and indirectly benefit coral communities by reducing algal competition. Consequently, conservation strategies aimed at restoring parrotfish populations are considered essential for maintaining resilient coral-dominated reef systems(Mumby et al., 2007; Li et al., 2014). The ecological importance of protecting parrotfish for endangered corals is substantial. Under normal conditions, parrotfish communities can maintain approximately 40% of coral reefs under consistent grazing pressure, whereas overfishing diminishes this capacity to merely 5%(Mumby et al., 2007; Li et al., 2014; Mcmanus et al., 2018; Nes et al., 2016). Sea urchins, when present in moderate numbers, function as even more effective herbivores than parrotfish. This was dramatically demonstrated in 1983 when mass sea urchin mortality led to a shift from coral dominance to algal dominance, leaving only the less efficient parrotfish as grazers. The critical transition dynamics between coral and algal states have been extensively investigated by numerous researchers(Mumby et al., 2007; Li et al., 2014; Nes et al., 2016; Mcmanus and Polsenberg, 2004; Hughes et al., 2007; Mccook et al., 2001; Dudgeon et al., 2010; Andersen et al., 2009). Research has established that coral-algae systems typically exhibit two distinct stable states: a coral-dominated condition and an algal-dominated condition(Mumby et al., 2007; Li et al., 2014; Mcmanus et al., 2018; Nes et al., 2016). This ecological bistability forms the conceptual foundation for our study. While recent work has explored more complex models incorporating recruitment seasonality and grazing effects(Mcmanus et al., 2018), our analysis focuses specifically on a simplified coral-algae interaction model(Mumby et al., 2007; Li et al., 2014) to investigate the critical factors determining reef ecosystem.

At low grazing intensities, where parrotfish consume macroalgae without distinguishing from algal turfs, coastal seabeds become covered by macroalgae, resulting in a macroalgal-dominant state. Conversely, high grazing intensities promote coral coverage, creating a coral-dominant state. When grazing pressure decreases below a critical threshold, coral populations decline while macroalgae proliferate, causing a shift from coral dominance to macroalgal dominance(Mumby et al., 2007; Li et al., 2014; Mcmanus et al., 2018). Within a specific range of grazing intensities, both macroalgal-dominant and coral-dominant states represent alternative stable state of the ecosystem(Mcmanus and Polsenberg, 2004; Hughes et al., 2007; Mccook et al., 2001; Dudgeon et al., 2010; Mumby et al., 2007; Li et al., 2014; Mcmanus et al., 2018).

State changes in complex ecological systems can be described through the mathematical frameworks of phase transitions or bifurcations. Nonlinear dynamical systems can exhibit various behaviors including steady states, periodic orbits, and chaotic dynamics. Much researches have predominantly focused on ecological stability at equilibrium points(Mumby et al., 2007; Li et al., 2014; Mcmanus et al., 2018). This approach typically examines the basins of attraction of these equilibria across different parameter values, thereby emphasizing local stability properties near equilibrium points (Scheffer et al., 2009). However, conducting global stability analysis of coral-algal systems presents significant challenges, and the relationship between systemwide dynamics and the behavior of individual components remains incompletely understood. In this study, we demonstrate how landscape-flux theory, derived from non-equilibrium statistical mechanics, provides an effective framework for analyzing the global stability properties of coral-algae ecosystems. We utilize a well-established coral-algal model as our primary case study (Mumby et al., 2007; Li et al., 2014).

Understanding how natural systems respond to human disturbances and identifying critical thresholds is essential for developing effective early warning systems for ecological transitions(Andersen et al., 2009; Bestelmeyer et al., 2013; Biggs et al., 2009). As ecosystems face increasing pressure from climate change, the ability to detect tipping points and anticipate critical transitions has become increasingly important(Lenton, 2011; Scheffer et al., 2015; Thompson and Sieber, 2011). Early warning signals play a crucial role in this process, helping us to understand when abrupt and significant changes might occur in complex ecological systems(Clements and Ozgul, 2018a; Contamin and Ellison, 2009; Drake and Griffen, 2010). Before reaching a critical point, ecosystems typically maintain a sustainable balance; however, once this threshold is crossed, the current stable state loses stability, triggering catastrophic shifts to alternative stable states(Dai et al., 2012, 2013; Scheffer et al., 2015).

Recent theoretical and empirical investigations have substantially advanced our understanding of ecological system instabilities(Carstensen et al., 2013; Dakos et al., 2012; Guttal and Jayaprakash, 2009; Kéfi et al., 2014). Critical slowing down (CSD) theory has emerged as a framework in this field and has been widely applied to predict warning signals from univariate time series data(Dakos et al., 2015; Lindegren et al., 2012; Veraart et al., 2012; Scheffer et al., 2001). This behavior occurs as a control parameter approaches a critical threshold value, causing system dynamics to decelerate while the current steady state becomes increasingly unstable(Berglund and Gentz, 2006; Hastings et al., 2018; Scheffer et al., 2012). Common indicators include increased variance, stronger autocorrelation, and longer return times following perturbations(Boettiger and Hastings, 2012; Dakos et al., 2012; Gsell et al., 2016).

Despite its theoretical promise, research has revealed significant limitations to CSD’s practical application. Time delays in ecological systems fundamentally alter the dynamical properties near critical transitions, potentially rendering CSD indicators unreliable or misleading(Guttal et al., 2013). This theoretical concern is substantiated by empirical evidence from natural systems, where comprehensive analyses of long-term data from aquatic ecosystems demonstrate that CSD indicators’ efficacy is considerably constrained by real-world complexity, with environmental stochasticity and multiple interacting stressors frequently obscuring warning signals(Gsell et al., 2016).

While recent advances have expanded CSD applications through refined statistical indicators(Boulton and Lenton, 2019; Bury et al., 2021a) and multivariate extensions(Weinans et al., 2019), significant limitations remain. Most notably, CSD often provides warnings only when systems are already near critical thresholds-frequently too late for effective intervention(Biggs et al., 2009; Boettiger and Hastings, 2013b; Ditlevsen and Johnsen, 2010). Additionally, while CSD performs reliably in one-dimensional systems, it struggles with complex multidimensional ecological dynamics involving feedback loops(Boerlijst et al., 2013; Hastings and Wysham, 2010; Weinans et al., 2019). These shortcomings, along with challenges such as false signal susceptibility(Boettiger et al., 2013b; Perretti and Munch, 2012) and extensive data requirements(Burthe et al., 2016), highlight the need for complementary approaches that can provide earlier warnings for complex ecological systems and overcome the limitations inherent in current methodologies(Boettiger and Hastings, 2013b; Clements and Ozgul, 2018a; Dakos et al., 2015).

There has also been considerable recent interest in early warning signals based on AI and machine learning methods(Grassia et al., 2021; Bury et al., 2021b). While these methods often show impressive results on simulated and training data, it remains to be seen how well they generalize to different physical systems and unseen datasets. Moreover, these methods have an inherent disadvantage in that the generated EWSs do not have a rigorous mathematical underpinning and are typically not as interpretable to practitioners working in the application area(George et al., 2023). Machine learning methods have also recently been used to predict critical transitions by using existing EWSs (including those based on CSD) as features in the models to leverage subject matter expertise and insights(Ma et al., 2018; Lassetter et al., 2021). This hybrid approach could be a promising direction for practical testing of EWSs, including our landscape-flux based indicators.

Early warning signals of critical transitions help us to anticipate and understand the likelihood of abrupt and significant changes in complex systems. Ecosystems can usually maintain a sustainable balance before reaching a critical point, but upon crossing the critical point, the current stable state can lose stability, triggering a catastrophic transition to a new stable state. Near the critical point, the mechanisms sustaining the functioning of the ecosystem can break down, resulting in a sudden loss of resilience and preventing recovery. It is crucial to detect signals of critical transition as early as possible to give enough time to avert a potential ecological crisis, and the search for early predictions of imminent structural changes has thus become the focus of intense research. Critical slowing down theory is among the most popular and well-known approaches, but its predictions are only valid near the bifurcation point. In coastal ecosystems specifically, the goal is to detect warning signals for transitions from valued states (such as coral-dominated reefs) to degraded states (like macroalgal-dominated reefs), as well as assess the likelihood of recovery transitions. Developing indicators that can predict both the impending degradation and potential recovery before critical transitions occur would have substantial practical significance for ecosystem management(Mumby et al., 2007; Li et al., 2014; Veraart et al., 2012; Scheffer et al., 2001).

Ecological systems are increasingly recognized as inherently multivariate complex systems, and the understanding of their high-dimensional dynamic behavior requires further development (Boettiger and Hastings, 2013a; Boettiger et al., 2013a; Nolting and Abbott, 2016; Lamothe et al., 2019; Abbott and Dakos, 2021). Conventional one-dimensional stochastic models may be missing crucial elements needed to describe behaviors generated by rotational curl forces among variables originated from high-dimensional system. Rather than using traditional ecological theories based on general equilibrium assumptions, we need to characterize ecological systems through non-equilibrium processes. Recent advances in nonequilibrium statistical mechanics offer valuable insights into understanding attractor state formation, stability, bifurcations, and phase transitions in both physical and biological systems(Xu et al., 2014b; Wang, 2015; Wang et al., 2008; Xu et al., 2012; Wang et al., 2011, 2010; Qian, 2006; Ge and Qian, 2010; Qian, 2009; Xu et al., 2021, 2023).

In this study, we propose early warning signals for detecting approaching phase transitions in complex ecological systems. First, we measure the entropy production rate, which quantifies the energy dissipation or “thermodynamic cost” required to maintain ecosystem states far from equilibrium. Second, we analyze the average flux, which represents the net directional movement or flow of the system through its state space, indicating the strength of forces driving ecological dynamics. Third, we calculate the difference between forward-time and backward-time cross-correlations between system variables, which measures time irreversibility-the statistical difference between observing the system’s behavior in normal versus reversed time sequences. Together, these metrics can detect changes in system dynamics before traditional indicators reveal impending critical transitions, potentially providing warning signals.

Our findings demonstrate that these nonequilibrium warning indicators exhibit turning points between bifurcations, enabling predictions for both upcoming transitions significantly earlier than traditional critical slowing down indicators, which only become apparent near bifurcation points. The potential-flux landscape theory presents a effective approaches for exploring the underlying mechanisms of ecological catastrophes and improving the ability to predict critical transitions.

## 2 Methods

### 2.1 Coral-Algal Model

We explore the dynamics of a typical coral-algae ecosystem model(Mumby et al., 2007; Li et al., 2014), whose schematic diagram is shown in Figure 1A. The ecosystem model contains three functional types: macroalgae (*X*), coral (*Y*), and algal turfs (*T*), entities that can be colonised by macroalgae, algal turfs or coral. Algal turf consists of communities of short, densely growing filamentous algae that form a “turf-like” covering layer on hard substrates in coral reefs, typically reaching only a few millimeters in height. Unlike macroalgae, these turfs develop a low, compact structure that creates distinctive microhabitats while serving as a entity in reef ecosystem dynamics(Mumby et al., 2007; Li et al., 2014; Mcmanus et al., 2018).

**Figure 1.**
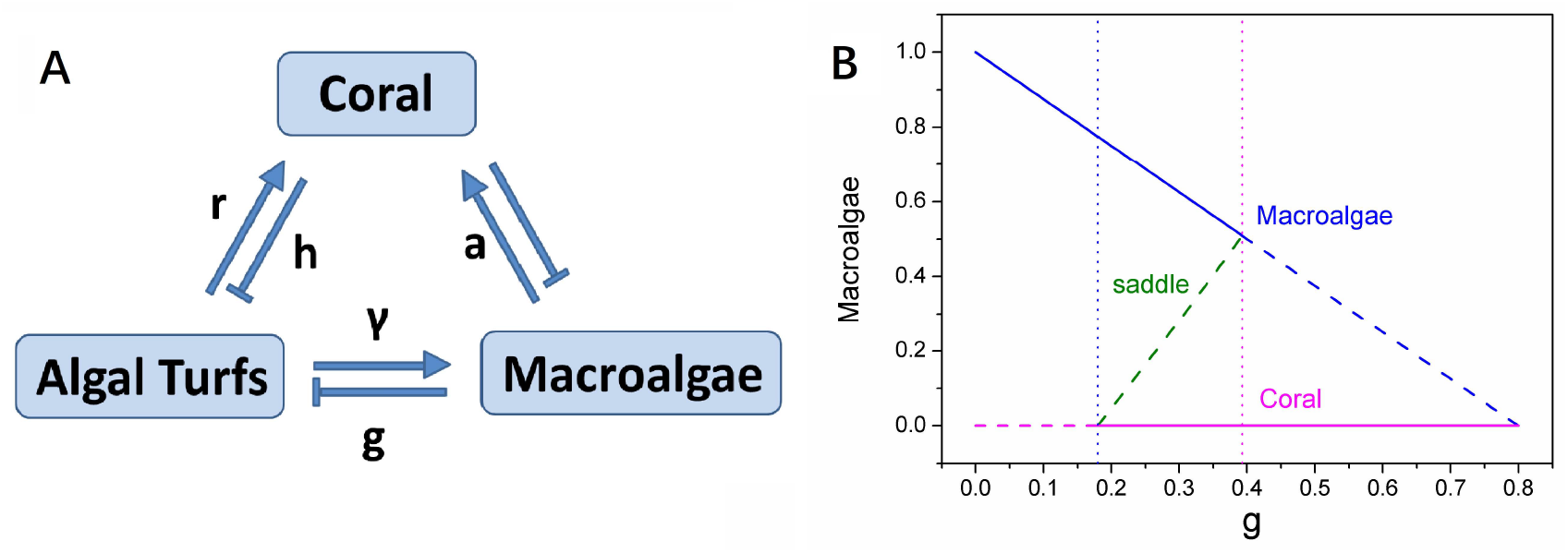
A: The schematic diagram for coral-algae model. B: The phase diagram versus grazing rate *g*.

We track the evolution of the proportions of space occupied by each functional type, effectively assuming that the system is spatially well-mixed, leading to a spatially implicit modelling framework(Mumby et al., 2007; Li et al., 2014; Mcmanus et al., 2018). This approach is appropriate for intermediate spatial scales where mixing processes (such as larval dispersal, water circulation, and mobile herbivore grazing) tend to homogenize local variations. The spatially implicit framework allows us to focus on ecosystem-level dynamics without the computational complexity of spatially resolved models. Corals recruit and overgrow algal turfs at rate *r*, while coral can be overgrown by macroalgae at rate *a*. Natural coral mortality occurs at rate *h* and we assume that space released by the death of the coral will be rapidly recolonized by algal turfs. Macroalgae colonizes algal turfs by covering them vegetatively at rate *γ*. Reef grazers, such as parrotfish, are assumed to consume macroalgae and algal turfs equally at rate *g*, and algal turfs arise when macroalgae are grazed. Thus the rate of algal turf production as a function of macroalgae is given by the proportion of grazing that affects macroalgae, i.e. *gX/*(*X* + *T*)(Mumby et al., 2007; Li et al., 2014; Mcmanus et al., 2018).

The coral-algae system can thus be described by the following set of nonlinear ordinary differential equations (ODEs):

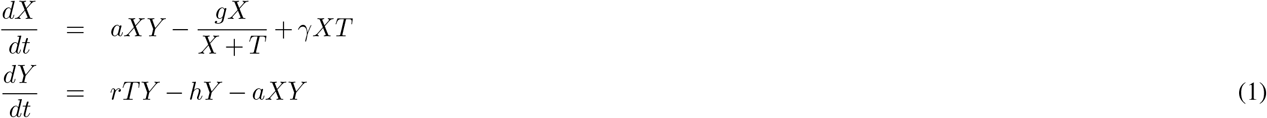

where *X* represents the proportion of space covered by macroalgae and *Y* represents the proportion of space covered by coral. *T* represents the proportion of algal turf cover and since we assume that all space (seabed) is completely covered by either macroalgae, coral or algal turfs, we have *X* + *Y* + *T* = 1, or *T* = 1 − *X* − *Y*. *g* is the grazing rate that parrotfish graze macroalgae without distinction from algal turfs range from 0 to 0.8. The parameter interpretations and their default values are given in Table I.

**Table 1.** Parameters interpretation and default values(Mumby et al., 2007; Li et al., 2014)

| Symbol | Ecological interpretation | Default value |
| --- | --- | --- |
| $a$ | the rate that corals are overgrown by macroalgae [$year^{-1}$] | 0.1 |
| $\gamma$ | the rate that macroalgae spread vegetatively over algal turfs [$year^{-1}$] | 0.8 |
| $r$ | the rate that corals recruit to and overgrow algal turfs [$year^{-1}$] | 1.0 |
| $h$ | the natural mortality rate of corals [$year^{-1}$] | 0.44 |
| $g$ | the rate at which herbivores consume macroalgae in the coral-algal model [$year^{-1}$] | |

Coral reef ecosystems can exhibit up to six distinct stable states: hard corals, turf algae, macroalgae, soft corals, coralimorpharians, and urchin barrens(Jouffray et al., 2015; Norström et al., 2009). While a more complex model incorporating all six states would better reflect ecological reality, we adopted a simplified two-state approach to facilitate analytical tractability while still capturing the fundamental bistable dynamics characteristic of critical transitions. This simplification enables us to clearly demonstrate the utility of our landscape-flux framework while maintaining mathematical accessibility. Additionally, our approach could potentially be extended to higher-dimensional systems with multiple stable states in future research, acknowledging both the limitations of our current model and opportunities for further development.

### 2.2 Landscape and flux theory for the coral-algae model

#### 2.2.1 The concept of landscape-flux theory

Landscape-flux theory provides a promising alternative framework for analyzing complex ecological systems and predicting critical transitions. This non-equilibrium statistical mechanics approach offers several distinct advantages over traditional methods. Foremost among these is its capacity to characterize global system stability through the construction of potential landscapes that quantify the relative stability of different states(Wang et al., 2008; Xu et al., 2014b, 2021). Unlike critical slowing down theory, landscape-flux theory effectively captures multidimensional system dynamics, including rotational forces (curl flux) as an additional driving force besides landscape gradient for the dynamics that are often overlooked in equilibrium-based analyses(Wang, 2015; Ge and Qian, 2010; Qian, 2006). This enables more comprehensive characterization of system behavior, particularly in complex ecological networks with multiple feedback mechanisms(Xu et al., 2023).

Another significant advantage is the theory’s ability to detect warning signals substantially earlier than bifurcation-proximity indicators(Wang et al., 2011, 2010). By quantifying both the potential landscape topography and the non-equilibrium flux, the approach provides mechanistic insights into transition drivers rather than merely phenomenological descriptions(Qian, 2009; Xu et al., 2012). The theory has been successfully applied to various complex systems(Wang, 2015; Fang et al., 2019), including gene regulatory networks(Wang et al., 2008), cell fate decisions(Wang et al., 2010; Xu et al., 2014b), and more recently, ecological regime shifts(Xu et al., 2021, 2023).

Despite its significant promise and advantages, landscape-flux theory presents certain challenges, particularly in its practical implementation. Its implementation requires sophisticated mathematical techniques and substantial computational resources(Wang, 2015; Fang et al., 2019). The approach demands comprehensive system knowledge for accurate model formulation and parameter estimation, which can be difficult to obtain for many ecological systems(Ge and Qian, 2010). Quantifying flux components in empirical systems poses challenges, often requiring high-resolution temporal data(Wang et al., 2011; Qian, 2009). There have not yet been any empirical studies combining the landscape flux theory and associated EWSs with data and it remains to be seen how successful the theory will be in practice. Nevertheless, the theory’s capacity to provide earlier warnings and deeper mechanistic understanding of ecological transitions makes it a valuable complement to existing approaches for analyzing complex ecosystems facing anthropogenic pressures and we hope that it can be empirically tested in the near future(Xu et al., 2021, 2023).

By adapting the potential landscape-flux framework to ecological dynamics, we bridge a critical gap between physical systems, where these methods originated, and complex biological systems characterized by nonlinear feedback and multiple stable states. Coral reef ecosystems represent an ideal test case for this theoretical extension due to documented evidence of alternative stable states, their sensitivity to environmental perturbations, and their growing vulnerability to climate change impacts. Our implementation demonstrates how landscape-flux theory can quantify stability of ecological systems under stochastic forcing, providing a mathematically rigorous foundation for early warning signals that complement existing early warning indicators for ecological systems(Clements and Ozgul, 2018b). This contrasts with some recent methods relying on AI and machine learning to produce indicators for transitions based on training on empirical data, but without a mathematical underpinning or basis through which to interpret the resulting indicators(George et al., 2023). Our work thus creates new opportunities for anticipating critical transitions in reef ecosystems, where traditional monitoring approaches often detect degradation only after substantial ecological changes have occurred. The framework’s ability to characterize global stability while accommodating environmental stochasticity makes it particularly suited to reef conservation, where identifying resilience thresholds and intervention windows is increasingly urgent for management and preservation efforts.

#### 2.2.2 Mathematics of Landscape and flux theory

The dynamics of the coral-algae model without noise or external fluctuations are characterized by a set of ordinary differential equations. In natural environments, however, coral-algae ecosystems are subject to diverse stochastic influences; internal stochasticity may emerge from variations in individual growth rates or grazing patterns, while external fluctuations may arise from processes such as ocean acidification, sedimentation, or other climate change-driven stressors(Scheffer et al., 2015; Carstensen et al., 2013). The deterministic model can be expressed in differential notation as *d***x** = **F**(**x**)*dt*, where vector **x** represents the ecosystem state and the driving force **F** encapsulates the interactions and transitions between coral, macroalgae, and algal turfs described in Figure 1A. To incorporate these various noise sources, we extend the model to:

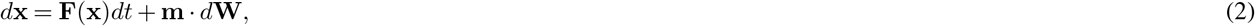

where **W**, coupled with matrix **m**, represents an independent Gaussian noise process(Gillespie, 1977; Wang, 2015; Swain et al., 2002). For analytical convenience, we define *D***G** = (1*/*2)(**m** · **m**^**T**^), where *D* is a constant representing the fluctuation scale and **G** is the diffusion matrix. Environmental disturbances such as temperature fluctuations, storm events, and nutrient pulses simultaneously affect coral, algae, and algal turfs, introducing correlations in the noise structure of natural reef systems. While our potential landscape-flux framework remains theoretically valid for systems with correlated noise, we have chosen to use a diagonal identity matrix for G to maintain analytical tractability. This simplification allows us to focus on the core dynamics while avoiding the substantial increase in mathematical complexity that would result from incorporating non-zero off-diagonal elements to represent correlated noise effects (Wang, 2015). Future extensions of this model could incorporate these more realistic noise structures to further refine predictions of reef dynamics under stochastic environmental forcing.

The probability of finding the coral-algae system in state **x** at time *t* is given by probability density function *P* (**x**, *t*), which evolves according to the Fokker-Planck equation(Van Kampen, 2007; Wang, 2015; Nicolis and Prigogine, 1977):

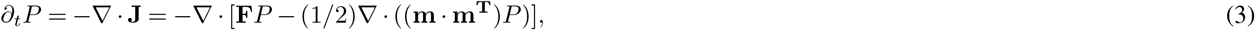

where **J** represents the probability flux through the system.

The steady-state probability distribution *P*_*ss*_(**x**) can be obtained by solving:

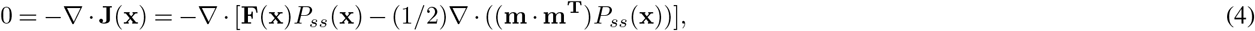

For equilibrium systems, we identify a “detailed balance solution” in which the flux **J** vanishes completely, signifying the absence of net energy transfer into or out of the system (Detailed discussion in Appendix A). In this case, *P*_*ss*_ ~ exp[−*U*](Gillespie, 1977; Van Kampen, 2007; Wang, 2015; Nicolis and Prigogine, 1977), where *U* represents the population-potential landscape. The driving force **F** can then be decomposed as:

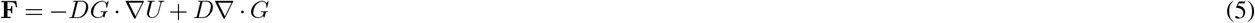

Thus, in equilibrium systems, **F** is determined entirely by the gradient of the potential landscape. We can calculate *P*_*ss*_ by solving equation or through experimental data collection, and subsequently derive the potential landscape using *U* = − ln *P*_*ss*_(Wang et al., 2008; Wang, 2015; Van Kampen, 2007).

For nonequilibrium systems, which better represent ecological reality(Hastings and Wysham, 2010; Weinans et al., 2019), the force decomposition becomes:

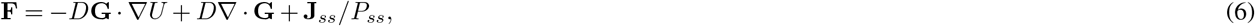

where **J**_*ss*_ denotes the non-zero steady-state probability flux, calculated as **J**_**ss**_ = **F***P*_*ss*_ − *D*∇ · (**G***P*_*ss*_). This flux satisfies ∇· **J**_*ss*_ = 0, indicating that **J**_*ss*_*/P*_*ss*_ represents a purely rotational force component. The potential gradient −*DG ·* ∇*U* drives the system toward stable states, while the divergence-free flux component generates rotational flow that facilitates transitions between alternative stable states(Wang et al., 2011; Xu et al., 2012). In nonequilibrium systems like coral reefs, both the potential landscape *U* and flux **J**_*ss*_ contribute to the system dynamics. Despite being conceptually derived from equilibrium theory, the potential landscape provides valuable insights into the global stability properties of nonequilibrium ecological systems(Xu et al., 2021, 2023), as we will demonstrate for the coral-algae model.

#### 2.2.3 Entropy production rate (EPR) and the average Flux (*F lux*_*av*_)

In nonequilibrium systems, the non-zero curl flux **J**_*ss*_ breaks detailed balance and provides a quantitative measurement of the system’s deviation from equilibrium(Xu et al., 2014b; Wang et al., 2008; Wang, 2015; Wang et al., 2011, 2010; Qian, 2006). This deviation metric is particularly valuable for investigating instabilities in the current state and detecting transitions to new stable states, making flux a critical component in developing early warning indicators for nonequilibrium ecological systems(Xu et al., 2021; Dakos et al., 2015). Fundamentally, flux provides a framework for analyzing nonequilibrium thermodynamics through entropy production. For the stochastic coral-algae model, the system entropy can be defined as *S*_*entropy*_ = − *∫ P* ln *Pd***x**. The temporal evolution of this entropy can be decomposed into two components: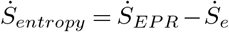, where 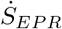 represents the entropy production rate (EPR) and 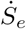 denotes the heat dissipation rate or environmental entropy change.

The entropy production rate is mathematically expressed as 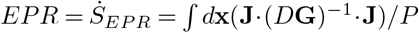(Qian, 2006; Wang et al., 2008; Zhang et al., 2012; Ge and Qian, 2010), while the heat dissipation rate is given by 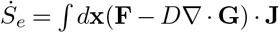.

The EPR is directly proportional to flux **J**, with larger flux values generating higher EPR values and consequently greater deviations from equilibrium(Ge and Qian, 2010; Qian, 2006). At steady state, a fundamental relationship emerges: the entropy production rate equals the heat dissipation rate(Ge and Qian, 2010; Qian, 2006; Wang et al., 2008; Zhang et al., 2012). In our analysis of the stochastic coral-algae model, we will utilize both the EPR and the average flux magnitude, defined as *Flux*_*av*_ = *∫*|**J**|*d***x**, to quantify the degree of nonequilibrium behavior and generate early warning signals for critical transitions(Wang et al., 2011; Xu et al., 2023).

#### 2.2.4 Time irreversibility: The average difference between forward and backward of cross-correlation

Time irreversibility in dynamical trajectories provides an effective method for quantifying nonequilibrium behavior in complex systems. We analyzed long-time trajectories of coral (*X*) and macroalgal (*Y*) cover simulated from the Langevin equation, focusing on noise-induced transitions between the *Macroalgae* and *Coral* attractors. The cross-correlation function forward in time is defined as *C*_*XY*_ (*τ*) = *(X*(0)*Y* (*τ*)), where *X* and *Y* represent time trajectories with interval *τ* (Qian and Elson, *2004; Zhang and Wang, 2018). Correspondingly*, 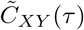 represents the cross-correlation function backward in time. The average difference between forward and backward cross-correlation, defined as 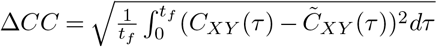, effectively quantifies time irreversibility. This measure captures the degree of nonequilibrium and flux strength through the system’s deviation from detailed balance(Qian and Elson, 2004; Zhang and Wang, 2018; Xu and Wang, 2020), offering a practical indicator of phase transitions directly observable from temporal trajectories.

#### 2.2.5 Escape time (The mean first passage time)

Ecological systems may transition from their current stable state to an alternative stable state due to stochastic fluctuations or external forces, effectively escaping their basin of attraction. The escape time between stable states provides a valuable quantitative measure for assessing global stability in coral reef ecosystems. By estimating the mean exit time from a basin of attraction (Arani et al., 2021; Wang et al., 2008; Xu et al., 2021, 2014a), we can better understand the likelihood of transitions between coral-dominated and algae-dominated states. Mean first passage time (MFPT)-the average time required for a stochastic process to first reach a specified threshold value-provides a robust metric for quantifying this phenomenon. MFPT effectively measures the kinetic speed or temporal characteristics of transitioning between states, offering natural indicators of a system’s propensity to depart from its current basin of attraction.

To investigate this behavior, we employ Langevin dynamics to simulate the stochastic coral-algae model and analyze the MFPT distribution between stable states. Our methodology begins by selecting one stable state as the initial condition, while designating a disc with radius *r*_0_ = 0.01 surrounding the alternative stable state as the target “state.” We then compile first passage time statistics from the initial to the final state, subsequently averaging across all simulations to determine the mean first passage time. We define *τ*_*CM*_ as the *MFPT* from the coral-dominated state to the macroalgae-dominated state, and conversely, *τ*_*MC*_ as the *MFPT* from the macroalgae-dominated state to the coral-dominated state.

#### 2.2.6 Lyapunov function for the coral-algae model under zero fluctuations

In dynamical systems theory, Lyapunov functions serve as powerful tools for stability analysis, enabling characterization of an attractor’s global stability beyond the limitations of local stability analysis(Wang, 2015; Fang et al., 2019). We discussed the differences between global stability and local stability detailed in SI. While no general method exists for constructing Lyapunov functions for complex nonlinear systems, we can utilize the steady-state probability distribution *P*_*ss*_ and the population potential *U* to investigate the global stability properties of the stochastic coral-algae model under finite fluctuations. Unfortunately, the population potential landscape *U* does not generally function as a Lyapunov function(Xu et al., 2014a; Zhang et al., 2012), in the small noise limit (*D →* 0^+^), the intrinsic potential landscape *ϕ*_0_ emerges as a viable Lyapunov function. We can compute *ϕ*_0_ by solving the Hamilton-Jacobi equation:

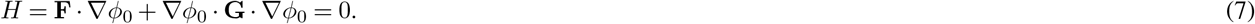

This equation results from expanding the population potential *U* in powers of noise level *D*, substituting this series into the Fokker-Planck equation, and truncating at order *D*^−1^ to obtain the equation for *ϕ*_0_(Wang, 2015; Xu et al., 2014a; Zhang et al., 2012).

To verify that *ϕ*_0_ functions as a Lyapunov function, we calculate:

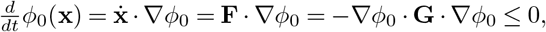

where the inequality holds when **G** is positive definite. This demonstrates that *ϕ*_0_(**x**) monotonically decreases along deterministic trajectories as *D →* 0, confirming its utility for quantifying global stability in the small noise regime. The intrinsic potential landscape *ϕ*_0_ relates to the steady-state probability and population potential landscape through *U* = −*lnP*_*ss*_ ~ *ϕ*_0_*/D* as *D →* 0.

In the zero fluctuation limit, the driving force **F** can be decomposed into gradient and curl components:

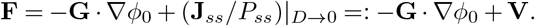

The first term, −**G** · ∇*ϕ*_0_, represents the gradient of the non-equilibrium intrinsic potential, while **V** = (**J**_*ss*_*/P*_*ss*_)_*D→*0_ defines the intrinsic steady-state flux velocity. The steady-state intrinsic flux term **J**_*ss*_|_*D→*0_ is divergence-free due to ∇ · **J** = 0. From the Hamilton-Jacobi equation, we derive (**J**_*ss*_*/P*_*ss*_)|_*D→*0_ · ∇*ϕ*_0_ = **V** · ∇*ϕ*_0_ = 0, establishing that the intrinsic potential gradient is perpendicular to the intrinsic flux in the zero fluctuation limit(Wang et al., 2011, 2010).

For the coral-algae model, calculating the intrinsic potential *ϕ*_0_ presents substantial difficulties due to the constrained state space (an isosceles triangle where 0 ≤ *X, Y* ≤ 1, 0 ≤ 1−*X* −*Y* ≤ 1). The intrinsic potential is challenging to compute from the Hamilton-Jacobi equation in a normalized triangular state space. These geometric constraints complicate the analytical solution of the Hamilton-Jacobi equation, requiring specialized mathematical approaches to capture the system’s dynamical properties within this bounded domain. We therefore expand the potential *U* (**x**) in the small diffusion limit as *U* (**x**) = *ϕ*_0_(**x**)*/D*+*ϕ*_1_(**x**)+ *O*(*D*^2^) and employ a linear fitting method to approximate *ϕ*_0_. By plotting diffusion coefficients *D* versus *DU* (specifically *D* ln *P*_*ss*_) using small *D* values, we determine *ϕ*_0_ from the slope of the resulting line(Zhang et al., 2012; Xu et al., 2021, 2023). Additional analyses presented in the Supplementary Information include non-equilibrium thermodynamics, entropy dynamics, energy and free energy characteristics under both zero-fluctuation and finite fluctuation conditions, and kinetic pathways between alternative stable states (Macroalgae and Coral) in the model system. We add a glossary of terms in Table S1.

## 3 Results

Applying the landscape-flux framework described above, we now examine the dynamics and stability properties of the coral-algal ecosystem model under both finite and zero fluctuation conditions. Throughout our analysis, we distinguish between two alternative stable states: the “*Macroalgae*” state, characterized by macroalgal dominance and low coral density or by macroalgal only, and the “*Coral*” state, defined by coral dominance and minimal macroalgal presence or by Coral only(Mumby et al., 2007; Scheffer et al., 2015). This bimodal pattern of community structure represents a classic example of alternative stable states in marine ecosystems, with critical implications for reef resilience and conservation(Hastings et al., 2018; Scheffer et al., 2001). By quantifying the potential landscape and probability flux patterns associated with these states, we aim to characterize global stability properties and develop early warning indicators for critical transitions between these alternative ecosystem.

In the model, parameter *g* represents the grazing rate of macroalgae by herbivorous fish and invertebrates, a crucial ecological process with well-documented real-world counterparts. This parameter directly connects mathematical modeling to measurable ecological dynamics that reef managers can monitor and potentially influence. Real-world factors affecting the grazing rate *g*, including overfishing of herbivores (decreasing *g*), establishment of marine protected areas (increasing *g*), disease outbreaks among key grazers like the 1983 Caribbean sea urchin die-off (reducing *g*), and predator-prey dynamics through trophic cascades. As *g* gradually decreases in natural systems, algae gain competitive advantage over corals, system resilience weakens, recovery becomes increasingly difficult after disturbances, and eventually, at the critical threshold, even minor herbivore loss can trigger a shift to algal dominance. This mechanism explains ecological transitions observed on reefs, where reduced herbivory caused coral-to-algae phase shifts matching our bifurcation analysis predictions.

Figure 1B illustrates the deterministic phase diagram of the coral-algae system as a function of the parrotfish grazing rate *g* (which acts on macroalgae without distinguishing from algal turfs). When 0 ≤ *g* < 0.1796, the system exhibits one unstable fixed point (the dashed Coral state) and one stable fixed point (the solid Macroalgae state), indicating macroalgal dominance. As grazing intensity increases to 0.1796 ≤ *g* < 0.3927, the system transitions to bistability, characterized by two stable fixed points-the solid Macroalgae state and the solid Coral state-separated by an unstable green saddle fixed point that serves as a threshold between the two stable regimes. This bistable configuration persists until *g* = 0.3927, beyond which (*g* > 0.39) only the coral-dominated fixed point remains stable, indicating a complete shift to coral dominance at higher grazing intensities. The diagram which denotes the noise induced transitions with parameter driven reveals a bistable region wherein two alternative stable states-*Macroalgae* and *Coral*-coexist across a specific range of grazing values. This bistable region is bounded by transcritical bifurcations, which occur precisely when one equilibrium solution enters or exits the ecologically feasible region of phase space (defined by 0 ≤ *X* + *Y* + *T* ≤ 1, 0 ≤ *X, Y, T* ≤ 1).

To characterize the global stability properties of this system, we solved the Fokker-Planck equation for the coral-algae model, yielding the steady-state probability distribution *P*_*ss*_ and consequently the population landscape via *U* = −*lnP*_*ss*_. Figure 2A presents three-dimensional visualizations of these population-potential landscapes under finite fluctuations (*D* = 0.0005). These landscapes reveal how system stability evolves with changing grazing pressure. At low grazing rates (0 ≤ *g* < 0.1796), the landscape exhibits a single stable state dominated by macroalgae (the *Macroalgae* state). As grazing intensity increases 0.1796 ≤ *g* < 0.3927, a bistable landscape emerges with local minima corresponding to both macroalgal and coral dominance. With further increases in grazing rate, the *Coral* state deepens while the *Macroalgae* state becomes increasingly shallow and eventually disappears (*g* > 0.39), as also conceptualized in Figure 1B. At sufficiently high grazing rates, macroalgae are effectively eliminated from the system, and the landscape exhibits a single deep basin corresponding to coral dominance.

**Figure 2.**
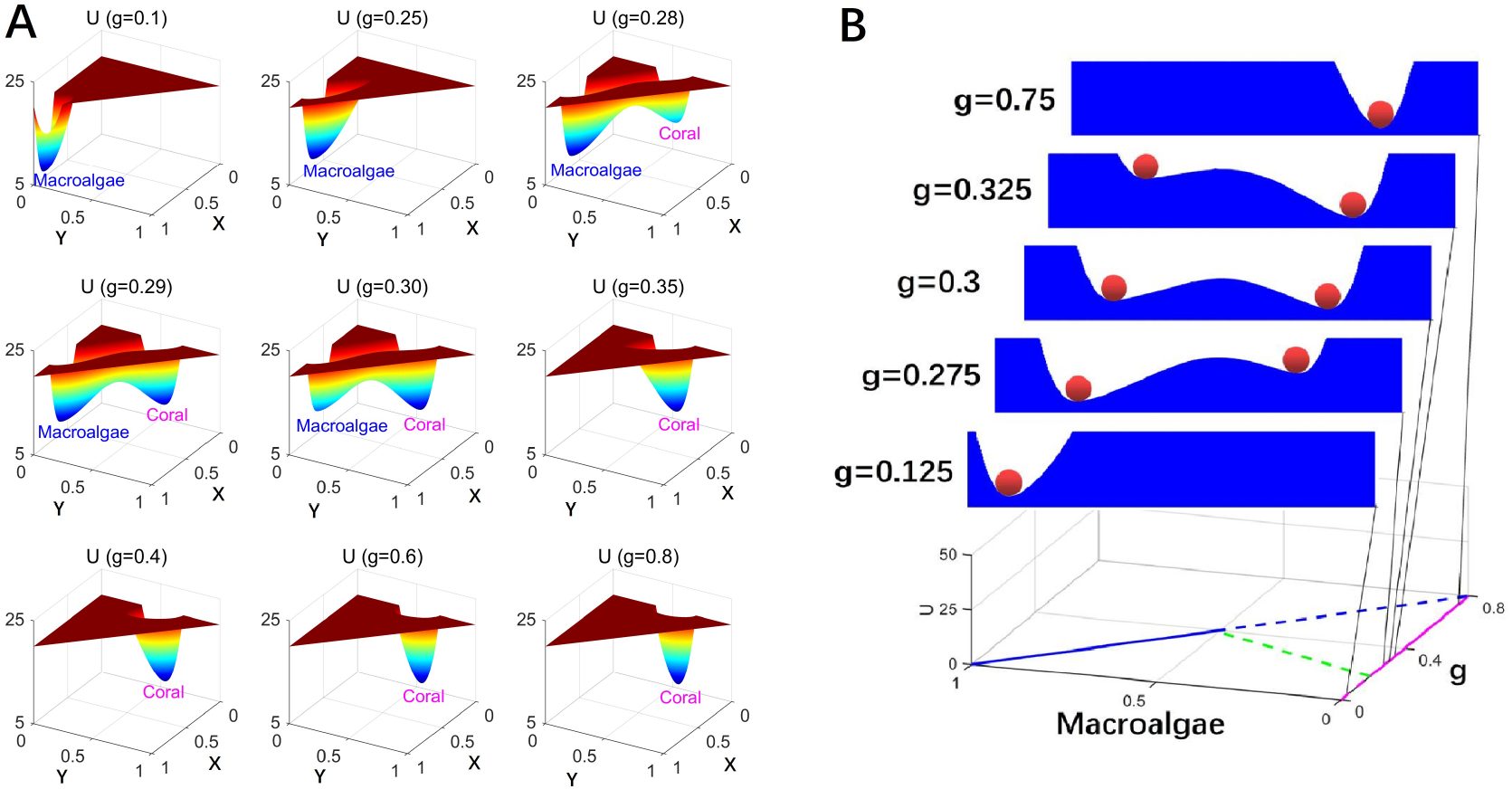
A: The population potential landscape *U* for the coral-algae model with finite fluctuation *D* = 5 × 10^4^. B: The population potential landscape *U* projected on *X*.

This progression of landscape topographies provides a comprehensive visualization of how grazing pressure drives transitions between alternative community states in coral-algae ecosystems(Scheffer et al., 2012; Hastings et al., 2018).

Natural ecosystems invariably experience disturbances and stochastic fluctuations. In systems characterized by alternative stable states, sufficiently intense fluctuations can propel the system from one stability basin through an unstable threshold, resulting in transition to an alternative stable configuration. Figure 2B illustrates this dynamic process through the classical “ball-in-the-valley” conceptual model(Scheffer et al., 1993; Scheffer, 2009), which visually represents the population potential landscape *U* projected onto coral cover (*X*) under different grazing intensity (*g*). This potential landscape is quantitatively derived from the steady-state probability distribution of the stochastic coral-algal model.

In this visualization, the ecosystem state is represented by a ball that naturally moves downhill and stabilizes in potential basins (valleys) that vary with grazing intensity. Each valley corresponds to an attraction basin in dynamical systems theory(Nolting and Abbott, 2016; Lamothe et al., 2019; Abbott and Dakos, 2021). Under small fluctuations, the system may temporarily deviate from equilibrium (the ball climbs partway up the slope) before returning to its steady state at the basin minimum. However, sufficiently large fluctuations can drive the system across the ridge (passing an unstable saddle point) into an alternative stability basin.

The landscape topography undergoes systematic transformations as grazing intensity increases: at *g* = 0.125, only the macroalgal valley (*M*) exists; at *g* = 0.275, both valleys exist but the coral valley (*C*) remains shallower than the macroalgal valley; at *g* = 0.3, both valleys attain similar depths, indicating comparable stability; at *g* = 0.325, the coral valley becomes deeper than the macroalgal valley; and finally, at *g* = 0.75, only the coral valley remains. This progression captures the grazingmediated shift from macroalgal to coral dominance in reef ecosystems.

Figure 3A-C demonstrates that the intrinsic potential landscapes calculated for the coral-algae model exhibit qualitatively similar patterns to the corresponding population potential landscapes across the grazing gradient, further validating the stability analysis approach. Figure 3D illustrates the intrinsic flux (purple arrows) and the negative gradient of the intrinsic potential landscape (white arrows) at grazing rate *g* = 0.29, clearly depicting their directional relationships in the vicinity of steady states. A striking feature of these vector fields is their orthogonality-the intrinsic fluxes are perpendicular to the negative gradients of the intrinsic potential landscape −∇*ϕ*_0_. This perpendicularity emerges from the mathematical relationship (**J**_*ss*_*/P*_*ss*_)|_*D→*0_ · ∇*ϕ*_0_ = **V** · ∇*ϕ*_0_ = 0, which is derived from the Hamilton-Jacobi equation under the zero fluctuation limit.

**Figure 3.**
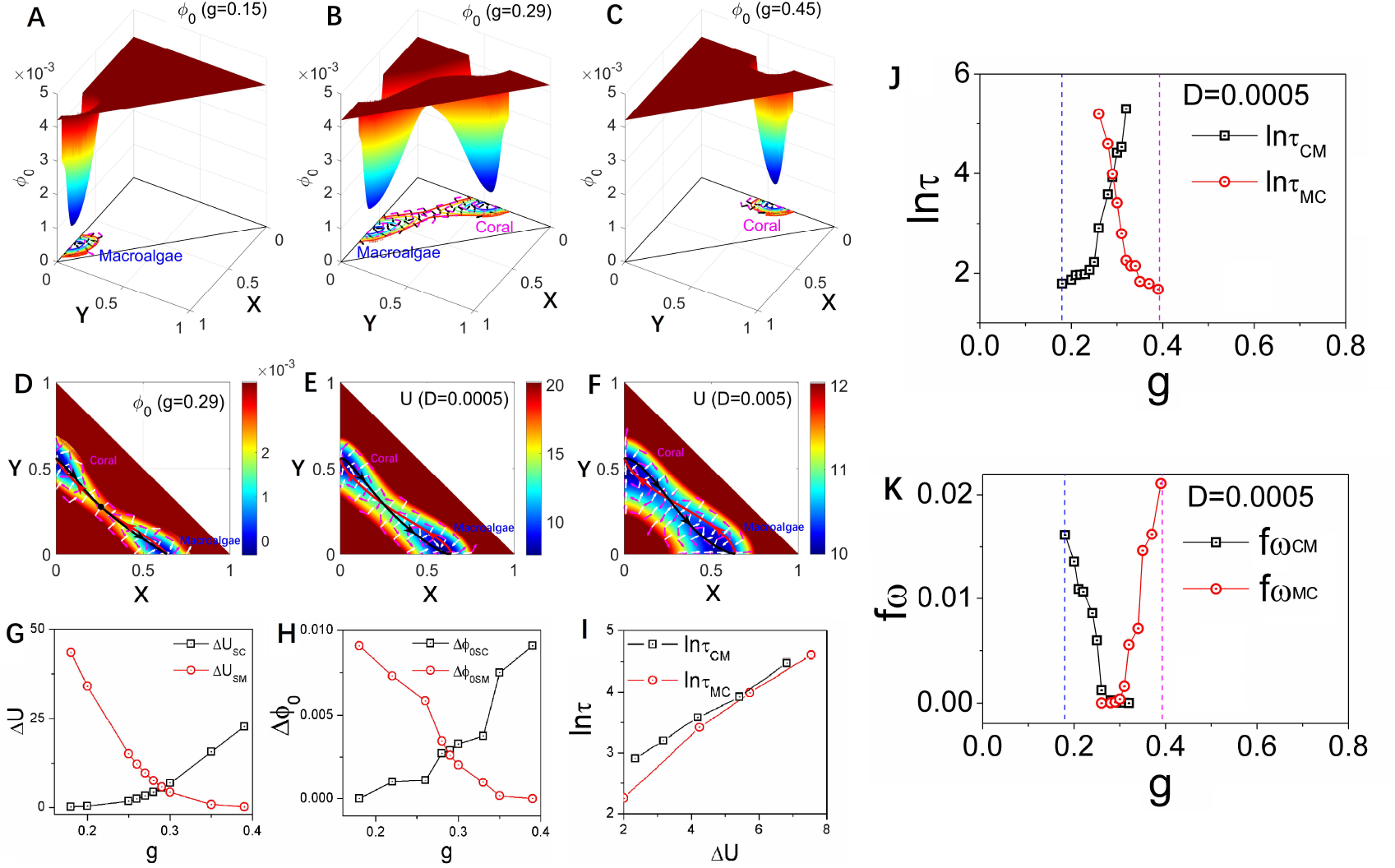
A-C: The intrinsic potential landscape with different *g* for the coral-algae model. D-F: The dominant intrinsic paths and fluxes on the intrinsic-potential landscape *ϕ*_0_ with zero fluctuation limit and grazing rate *g* = 0.29 (D). The dominant population paths and fluxes on the population-potential landscape *U* with the diffusion coefficient *D* = 0.0005 (E), *D* = 0.005 (F). The red lines represent the dominant paths from the *Macroaglae* state to *Coral* state. The black lines represent the dominant paths from the *Coral* state to *Macroaglae* state. The white arrows represent the steady-state probability fluxes. G: The population barrier heights versus parameter *g*. H: The intrinsic barrier heights versus parameter *g*. I:The population barrier heights versus the mean first passage time. Population barrier height Δ*U*_*SC*_ = *U*_*S*_ *– U*_*C*_ and intrinsic barrier height Δ*ϕ*_0*SC*_ = *ϕ*_0*S*_ −*ϕ*_0*C*_, Δ*U*_*SM*_ = *U*_*S*_ −*U*_*M*_ and Δ*ϕ*_0*SM*_ = *ϕ*_0*S*_ −*ϕ*_0*M*_ and *τ*_*CM*_ represents the mean first passage time from state *Coral* to state *Macroalgae* and *τ*_*MC*_ represents the mean first passage time from state *Macroalgae* to state *Coral*. J: The logarithm of MFPT versus *g*. K: The frequency of the flickering *fω* versus grazing rate *g*.

Figures 3E and F display the flux (purple arrows) and negative gradient of the population potential landscape (white arrows) superimposed on the landscape for different fluctuation intensities: *D* = 0.0005 (E) and *D* = 0.005 (F). The circulating fluxes around the stable states enhance communication between the *Macroalgae* and *Coral* states. These visualizations effectively demonstrate how the driving forces of the coral-algal system can be decomposed into complementary components: **F** = −*D ·* ∇*U* + **J**_*ss*_*/P*_*ss*_ + *D*∇· **G** for finite fluctuations and **F** = −**G** · ∇*ϕ*_0_ +(**J**_*ss*_*/P*_*ss*_)|_*D→*0_ = −**G** · ∇*ϕ*_0_ + **V** for the zero fluctuation limit. The substantial difference in magnitude of color bar units reflects the fundamentally different metrics being visualized: Figures 3D represents the intrinsic potential landscape derived from the Hamilton-Jacobi equation with zero limit fluctuations, whereas Figures 3E and Figures 3F show the population potential landscape from the Fokker-Planck equation with finite fluctuations. These inherent mathematical differences naturally produce different numerical ranges.

Figure 3D,E,F further reveals the dominant transition pathways between alternative stable states. Red lines represent the dominant paths from the *Macroalgae* state to the *Coral* state, while black lines indicate dominant paths in the reverse direction-shown on both the intrinsic potential landscape *ϕ*_0_ under zero fluctuations (D) and the population-potential landscape *U* under finite fluctuations (E and F). The purple arrow fluxes in Figure 3D guide these dominant paths under zero fluctuations, causing them to deviate from the steepest descent paths and diverge from each other as they pass through the saddle point-a deviation from equilibrium systems where zero flux would result in convergent paths. Similarly, under finite fluctuations (Figure 3EF), the dominant population paths guided by the purple arrow fluxes also deviate from steepest descent trajectories. This analysis reveals a fundamental feature of nonequilibrium systems: path irreversibility. The dominant paths from *Macroalgae* to *Coral* differ significantly from those in the reverse direction. This irreversibility stems from the nonequilibrium rotational flux, which creates spiral-shaped currents around stability basins. Interestingly, the dominant paths under zero fluctuation limit appear closer to each other compared to those under finite fluctuations, though they remain distinct due to the non-zero intrinsic flux. These spiral flux patterns represent the dynamical signature of nonequilibrium behavior in the coral-algal ecosystem(Wang et al., 2011; Xu et al., 2012; Zhang et al., 2012).

Figure 3G and H illustrate how barrier heights in both population-potential and intrinsic potential landscapes vary with grazing rate *g*. As *g* increases, the coral-algal system transitions from *Macroalgae* state dominance to *Coral* state dominance. This transition is reflected in the changing barrier heights: population barrier height Δ*U*_*SC*_ = *U*_*S*_ − *U*_*C*_ and intrinsic barrier height Δ*ϕ*_0*SC*_ = *ϕ*_0*S*_ − *ϕ*_0*C*_ increase with higher grazing rates, while Δ*U*_*SM*_ = *U*_*S*_ − *U*_*M*_ and Δ*ϕ*_0*SM*_ = *ϕ*_0*S*_ − *ϕ*_0*M*_ decrease. Here, *U*_*S*_ and *ϕ*_0*S*_ represent the potential values at the saddle point between alternative states, while *U*_*M*_, *ϕ*_0*M*_, *U*_*C*_, and *ϕ*_0*C*_ denote the minimum potential values in the *Macroalgae* and *Coral* states, respectively. These patterns demonstrate that elevated parrotfish grazing progressively destabilizes the *Macroalgae* state while enhancing the stability of the *Coral* state. The deeper attraction basin with higher barrier heights creates greater resistance to state transitions. Notably, both population and intrinsic barrier heights display nearly identical trends as *g* increases.

Figure 3J presents the Mean First Passage Time (MFPT), which quantifies the average time required for a stochastic process to first reach a specified state. The behavior of the mean first passage time (MFPT) as it is represented in logarithmic form, specifically shows an increase in ln *τ*_*CM*_ and a decrease in ln *τ*_*MC*_ as the parameter *g* increases. This trend indicates that it takes more time to exit the Coral state while it requires less time to transition out of the Macroalgae state as *g* rises. Consequently, the MFPT can effectively characterize the transition from the Macroalgae state to the Coral state with increasing *g*, providing a measurable indicator of this critical transition.

Figure 3I illustrates the logarithmic MFPT plotted against population barrier heights reveals a positive correlation-both ln *τ*_*CM*_ and ln *τ*_*MC*_ increase with barrier height, approximating a relationship of *τ* ~ exp(Δ*U*). This exponential relationship indicates that escape time dramatically lengthens as barrier height increases, directly linking transition kinetics to landscape topography. Specifically, a higher barrier height or deeper valley results in a longer time required to escape from that valley. This correlation suggests that the population-potential landscape topography is closely related to the kinetic speed of state switching, thereby influencing the communication capability for the global stability of the system.

The flickering frequency quantifies the number of state transitions per unit time. Specifically, *f*_*ωCM*_ represents the frequency of transitions from the *Coral* state to the *Macroalgae* state per unit time. In Figure 3K, we illustrate the frequency of transitions from *Macroalgae* to *Coral* (*f*_*ωMC*_) with fluctuation strength *D* = 5 × 10^−4^. Our results demonstrate that *f*_*ωMC*_ increases dramatically as *g* increases. This phenomenon can be explained by the decreasing stability of the *Macroalgae* state’s basin of attraction, which becomes shallower as *g* increases. Consequently, the system exhibits a higher probability of transitioning to the *Coral* state. Previous research has established flickering frequency as an effective early warning signal for critical transitions Scheffer et al. (2012, 2009). The tipping points identified through flickering frequency occur near the bifurcation point in the coral-algae model, where the *Macroalgae* state becomes unstable (flat potential) while the *Coral* state becomes dominant. Flickering frequency indicates that the *Macroalgae* state loses resilience, characterized by a diminishing basin of attraction in the potential landscape, while the *Coral* state gains dominance. It is important to note that actual transitions may occur considerably earlier than this bifurcation point due to larger environmental fluctuations.

While the effectiveness of critical slowing down as early warning indicators is under low noise conditions, a critical question remains regarding their robustness under more realistic, higher noise scenarios. This consideration is particularly important given that traditional critical slowing down indicators are known to perform poorly with increased stochastic fluctuations(Hastings and Wysham, 2010). We conducted analyses systematically varying the noise magnitude from *D* = 1 × 10^−5^ to *D* = 1 × 10^−2^ (Figure 4A-J). Figure 4 demonstrates how population entropy production rate *EPR* (A-E) and average flux *Flux*_*av*_ (F-J) vary with grazing rate under increasing finite fluctuations. Our findings reveal that both EPR and average flux (*F lux*_*av*_) maintain relatively robust performance as early warning signals up to *D* = 1 × 10^−2^, beyond which signal reliability begins to deteriorate significantly. This represents a substantial improvement over conventional critical slowing down indicators, which typically lose effectiveness at high noise levels. The relative noise robustness of our framework likely stems from the fact that our indicators directly quantify system-wide properties reflecting global stability, rather than local temporal patterns that become increasingly masked by higher noise.

**Figure 4.**
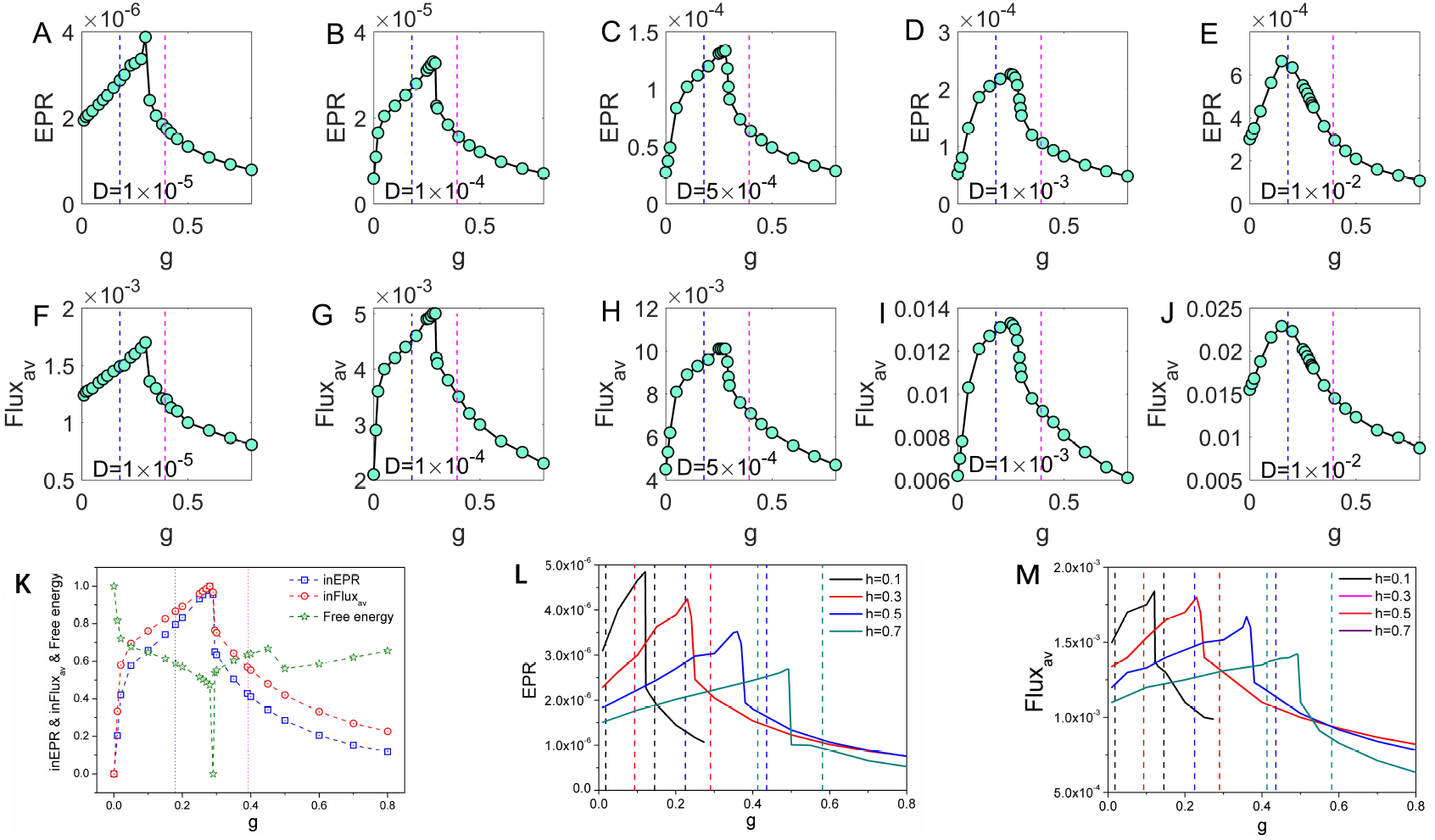
The population entropy production rate (A-E) and the population average flux (F-J) versus grazing rate *g* with increasing *D*. K: The intrinsic entropy production rate, the population average flux and the free energy versus grazing rate *g* for the coral-algae model (parameters are set in Table I). L-M: The *EPR* and *Flux*_*av*_ versus grazing rate *g* with different natural mortality rate of corals *h* for the coral-algal model. The dashed lines represent the locations of the transcritical points for each value of *h* with the same color for the *EPR* lines.

Figure 4K displays the intrinsic entropy production rate *inEPR* and intrinsic average flux *inFlux*_*av*_ against *g*. These two metrics exhibit a similar pattern-initially increasing and subsequently decreasing with higher grazing rates, with pronounced peaks occurring between the two transcritical bifurcations shown in Figure 4K. These peaks coincide with the critical transition region from *Macroalgae* to *Coral* dominance. Additionally, Figure 4K reveals that intrinsic free energy reaches a minimum near the peaks of *inEPR* and *inFlux*_*av*_. Collectively, these findings suggest that *EPR, Flux*_*av*_, *inEPR, inFlux*_*av*_, and intrinsic free energy can serve as effective indicators for detecting phase transitions and bifurcations in coral-algal systems(Xu et al., 2021, 2023).

To calculate state-specific time irreversibility measures, we employed relatively small diffusion coefficients to prevent spontaneous transitions between alternative stable states. This approach allowed us to collect sufficient stochastic simulation data while the system remained within either the *Macroalgae* or *Coral* state. We denote the resulting irreversibility measures as Δ*CCM* (for trajectories within the *Macroalgae* state) and Δ*CCC* (for trajectories within the *Coral* state). Notably, once a system transitions to an alternative state, the pre-transition irreversibility measure can no longer predict the transition that has already occurred.

Figure 5A illustrates how both Δ*CCM* and Δ*CCC* exhibit pronounced peaks between the two transcritical bifurcations under small fluctuations (*D* = 1 × 10^−5^) with parameter *h* = 0.44. Figures 5B display the derivatives of these measures: *k*_Δ*CCM*_ (the slope of Δ*CCM*) for the *Macroalgae* state, alongside *k*_Δ*CCC*_ (the slope of Δ*CCC*) for the *Coral* state. We fitted an exponential function to the simulation data to calculate these derivatives. The derivative measures exhibit clear inflection points, indicating significant changes as the system approaches bifurcation points. These characteristic patterns in irreversibility measures and their derivatives demonstrate their potential as early warning signals for critical transitions in coral-algae ecosystems.

**Figure 5.**
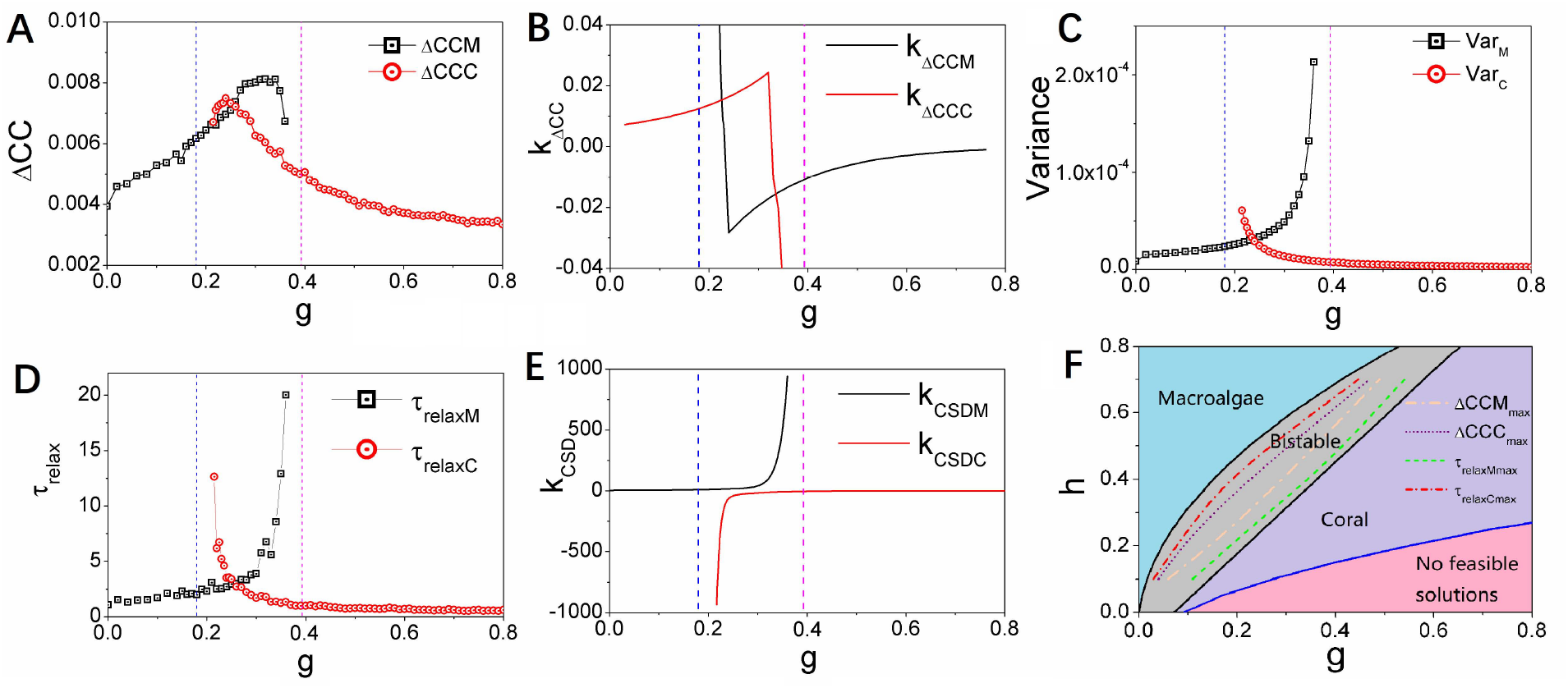
A: The average difference of the cross correlations forward and backward in time Δ*CCM* and Δ*CCC* versus *g*. B: *k*_Δ*CCM*_ (the slope of Δ*CCM*) and *k*_Δ*CCC*_ (the slope of Δ*CCC*)versus *g*. C: The variance *V ar*_*M*_ and *V ar*_*C*_ versus grazing rate *g*. D: The relaxation time *τ*_*relaxM*_ and *τ*_*relaxC*_ versus grazing rate *g*. E: *k*_*CSDM*_ (the slope of the relaxation time *τ*_*relaxM*_) and *k*_*CSDC*_ (the slope of the relaxation time *τ*_*relaxC*_) versus grazing rate *g*. F: The two-dimensional phase diagram of the natural mortality rate of corals *h* versus grazing rate *g* for the coral-algal model. Δ*CCM*_*max*_ represents the maximum of Δ*CCM*, Δ*CCC*_*max*_ represents the maximum of Δ*CCC. τ*_*relaxMmax*_ represents the coordinate position of the sharp rise of *τ*_*relaxM*_, *τ*_*relaxCmax*_ represents the coordinate position of the sharp rise of *τ*_*relaxC*_. (*D* = 1.0 × 10^−5^, h=0.44)

Figure 5C illustrates the relationship between variance and grazing rate *g*. Specifically, as the grazing rate *g* increases: the variance of the Macroalgae state (*V ar*_*M*_) shows a clear increasing pattern, while simultaneously, the variance of the Coral state (*V ar*_*C*_) exhibits a decreasing trend. This divergent behavior in variances provides important insights into the system’s stability characteristics. The increasing variance in the Macroalgae state (*V ar*_*M*_) indicates growing instability and fluctuations in this state as grazing pressure intensifies. Conversely, the decreasing variance in the Coral state (*V ar*_*C*_) signifies that this state becomes more stable and resilient with increasing grazing pressure. These variance patterns serve as quantitative early warning indicators of the shifting stability landscape in the coral-algae system and help identify the approach toward critical transition points in this ecological model.

Critical slowing down emerges as ecosystems approach bifurcation points during gradual environmental changes. When a system within a stable state experiences external disturbance, it eventually returns to its original equilibrium after a characteristic period known as the relaxation time(Scheffer et al., 2009). This relaxation time represents the system’s adaptive response to environmental perturbations. When varying grazing rate *g*, bifurcations can be approached from either increasing or decreasing directions. Critical slowing down effectively identifies the left bifurcation (where *Macroalgae* becomes dominant and the *Coral* state flattens) when *g* decreases, and the right bifurcation (where *Coral* becomes dominant and the *Macroalgae* state flattens) when *g* increases. Figure 5D illustrates this phenomenon in the coral-algal model: the relaxation time *τ*_*relaxM*_ for the *Macroalgae* state increases sharply when approaching the right transcritical bifurcation point with increasing *g*, while the relaxation time *τ*_*relaxC*_ for the *Coral* state similarly increases when approaching the left transcritical bifurcation with decreasing *g*.

Figure 5E displays the derivatives of these relaxation times: *k*_*CSDM*_ (slope of *τ*_*relaxM*_), *k*_*CSDC*_ (slope of *τ*_*relaxC*_) plotted against grazing rate *g*. These slopes, particularly *k*_Δ*CCC*_ and *kk*_Δ*CCC*_, exhibit sharp increases as the system approaches bifurcation points, confirming that relaxation time lengthens near critical transitions. However, the analysis reveals a crucial advantage of nonequilibrium warning indicators (flux, entropy production rate, time irreversibility) over traditional critical slowing down indicators-they provide substantially earlier predictions of impending bifurcations. For instance, Figure 5A shows that peaks in time irreversibility measures (Δ*CCC* and Δ*CCM*) occur within the bistable zone, whereas peaks in relaxation times (Figure 5D) appear only at the immediate vicinity of bifurcation points.

The nonequilibrium measures-flux magnitude, entropy production rate, intrinsic free energy, and time irreversibility-collectively provide early warning signals that precede predictions from conventional methods. In the coral-algae model, these nonequilibrium indicators predict the transition from *Macroalgae* dominance to *Coral* dominance midway through the bistable region, rather than near the critical threshold at *g* = 0.3927 where the *Coral* state becomes dominant. This represents a significantly earlier warning than previously reported approaches(Veraart et al., 2012; Scheffer et al., 2001, 2012).

Our nonequilibrium warning indicators-flux, entropy generation rate, time irreversibility from cross-correlation analysis, and non-equilibrium free energy-consistently exhibit critical transitions between the two transcritical bifurcations in the coral-algal model. These indicators provide substantially earlier warnings compared to traditional critical slowing down signals. From the perspective of a system currently in the *Macroalgae* state, our nonequilibrium signals anticipate the right bifurcation (where *Macroalgae* becomes unstable while *Coral* becomes dominant) well before critical slowing down indicators detect this transition. Similarly, from the perspective of a system in the *Coral* state, our indicators predict the left bifurcation (where *Coral* becomes unstable while *Macroalgae* becomes dominant) earlier than critical slowing down. This positioning of nonequilibrium indicator turning points in the middle of the bistable region enables prediction of both bifurcations with considerable advance warning.

Critical slowing down indicators suffer from a fundamental limitation-they invariably miss one bifurcation in each parameter direction. For instance, as grazing rate *g* increases toward the right bifurcation, critical slowing down fails to detect the left transcritical bifurcation where the *Macroalgae* state dominates and the *Coral* state first appears as a shallow attractor. This occurs because critical slowing down only manifests when the landscape around the current attractor flattens near a bifurcation point. When approaching the right bifurcation, the system’s current *Macroalgae* state becomes flat, producing critical slowing down. However, near the left bifurcation with increasing *g*, the *Macroalgae* state remains dominant with a non-flat landscape, preventing critical slowing down from emerging. Consequently, critical slowing down cannot predict left bifurcations when in a *Macroalgae*-dominated state with increasing *g*, nor right bifurcations when in a *Coral*-dominated state with decreasing *g*.

Figure 4L, M display entropy production rate (*EPR*) and average flux (*F lux*_*av*_) plotted against grazing rate *g* across different coral natural mortality rates *h*. Every data curve exhibits a pronounced peak within its corresponding bistable region, confirming that both *EPR* and *Flux*_*av*_ effectively indicate phase transitions in coral-algal systems. Our nonequilibrium early warning signals emerge midway between bifurcations, providing much earlier predictions in both parameter directions compared to critical slowing down indicators that appear only near specific bifurcations. This bidirectional predictive capacity represents a significant advantage of our approach, as illustrated in Figures 4.

Critical slowing down has been widely used in models with saddle-node bifurcations. In our case, because the stable solutions leave the feasible region exactly at the point at which transcritical bifurcations occur, we effectively have the same qualitative dynamics that occur in models with saddle-node bifurcations. In particular, there is one stable and one unstable solution approaching the bifurcation and both solutions disappear after the bifurcation occurs. So far, most studies have been focused on effective one-dimensional methods, the results of which can often be applied to effective equilibrium systems where global stability can be quantified by landscape alone, without considering the key non-equilibrium ingradient component, i.e., flux(Veraart et al., 2012; Scheffer et al., 2001, 2012, 2009). Our fully vectorized high dimensional formulation of the potential-flux landscape can quantify the non-equilibrium by the non-zero curl flux, which can lead to a much richer complex dynamics with detailed balance breaking. In contrast, the equilibrium dynamics are determined entirely by the gradient of the potential landscape. Curl fluxes that break the detailed equilibrium play an important role in driving the dynamics of the non-equilibrium system.

Figure 5F presents a two-parameter phase diagram illustrating the relationship between coral natural mortality rate *h* and grazing rate *g*. Parameter *h* in our model represents coral mortality rate, encompassing the cumulative effects of diverse environmental stressors affecting reefs globally. These include rising sea temperatures that trigger coral bleaching events (significantly increasing h during thermal anomalies), ocean acidification that reduces calcification rates and weakens coral skeletons (gradually elevating *h*), pollution, sedimentation, and coastal development that impose direct physiological stress, and the increasing frequency and severity of coral diseases worldwide that directly contribute to higher h values. These real-world stressors operate across different temporal scales-from acute (bleaching events) to chronic (acidification)-which aligns with our analysis of how gradual versus rapid parameter shifts influence system dynamics and stability. The diagram features four distinct regions: a blue region where only the *Macroalgae* state is stable; a grey bistable region where both *Macroalgae* and *Coral* states are stable; a purple region where only the *Coral* state is stable; and a pink region without feasible stable states. These regions are delineated by bifurcation curves (black for transcritical points, blue for saddle node points). Figure S1 provides additional phase diagrams for different *h* values.

Figure S2 demonstrate how time irreversibility metrics capture approaching bifurcations which is a noise induced transitions versus parameter grazing rate *g* driven in the coral-algal model for increasing *h*. Comparing the positions of maximum values and sharp rises in these indicators reveals a crucial temporal advantage of time irreversibility measures over critical slowing down indicators. Within the bistable region shown in Figure S2, Δ*CCM*_*max*_ (position of maximum Δ*CCM*) occurs significantly earlier than *τ*_*relaxMmax*_ (position where *τ*_*relaxM*_ sharply rises, defined as where *kk >* 1 × 10^4^) as gradual parameter changes. Similarly, Δ*CCC*_*max*_ (position of maximum Δ*CCC*) appears much earlier than *τ*_*relaxCmax*_ (position where *τ*_*relaxC*_ sharply rises). The *τ*_*relaxMmax*_ line lies considerably closer to the right bifurcation boundary than the Δ*CCM*_*max*_ line, while the *τ*_*relaxCmax*_ line lies much nearer to the left bifurcation boundary than the Δ*CCC*_*max*_ line. These spatial relationships consistently demonstrate that time irreversibility measures (Δ*CC*) provide substantially earlier warning signals than relaxation time (*τ*_*relax*_) indicators from CSD theory as gradual parameter *g* changes with fluctuations, confirming their efficacy as early warning signals for critical transitions in coral-algae ecosystems.

Figure 6 A and D illustrate the average differences in cross-correlations over time, represented as Δ*CCM* and Δ*CCC*, plotted against the grazing rate *g*. These values provide insight into the dynamics of the coral-algae system, revealing how the interaction strength between states varies as grazing pressure changes. Figure 6 B and E display the relaxation times, *τ*_*relaxM*_ and *τ*_*relaxC*_, in relation to the grazing rate *g*. The relaxation time quantifies how quickly the system responds to perturbations, serving as a crucial indicator of the stability of the Macroalgae and Coral states under varying conditions. Figure 6 C and F present the variances *V ar*_*M*_ and *V ar*_*C*_ as functions of the grazing rate *g*. These variances reflect the degree of fluctuations within each state, highlighting how stability is affected as grazing pressure increases. It is noteworthy that for Figure 6, the fluctuation strength is set at *D* = 5.0 × 10^−4^, while for Figures D-F, the fluctuation strength increases to *D* = 1.0 × 10^−3^, with a fixed height parameter of *h* = 0.44. This variation in *D* is expected to have a significant impact on the observed relationships, further illustrating the delicate balance between grazing intensity and the stability of the coral-algae ecosystem. We observe that while increasing noise can also serve as an indicator for predicting state transitions, its predictive effectiveness diminishes relative to the performance observed at lower noise levels.

**Figure 6.**
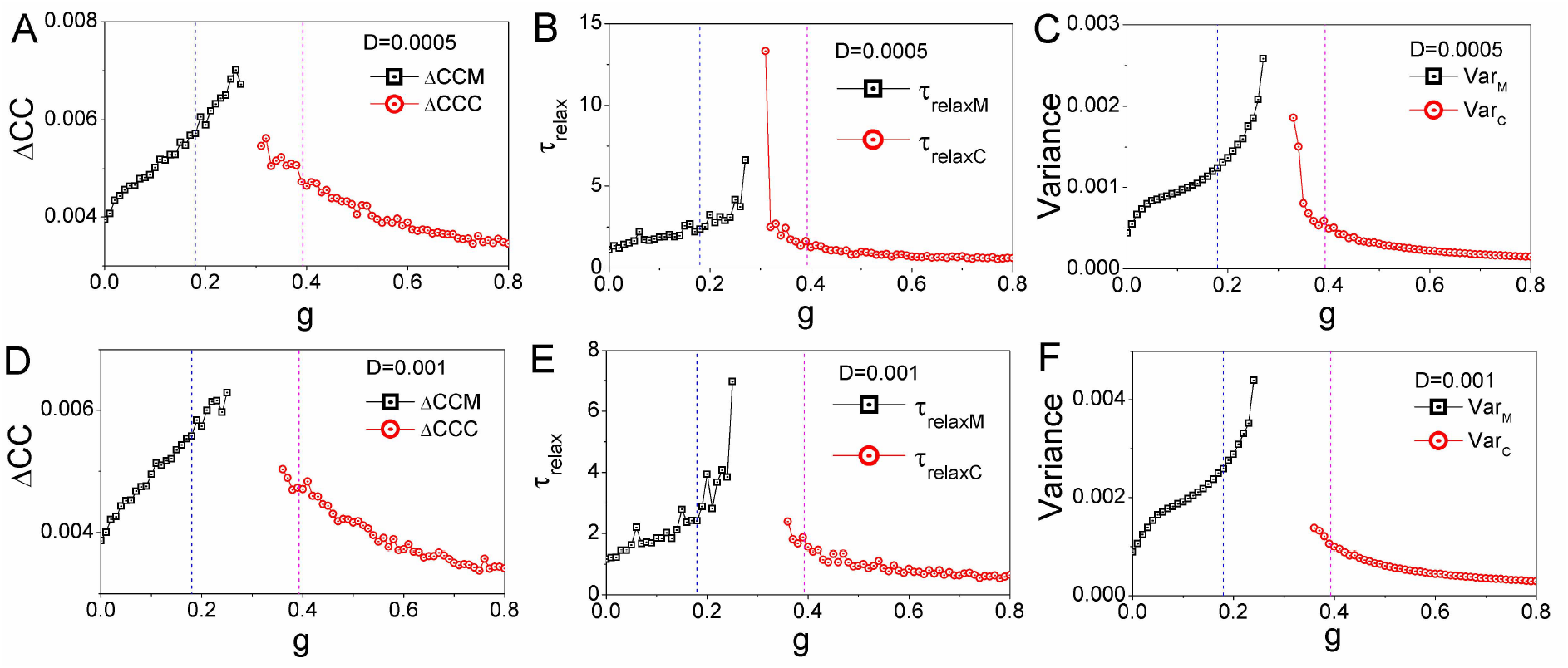
A,D: The average difference of the cross correlations forward and backward in time Δ*CCM* and Δ*CCC* versus *g*. B,E: The relaxation time *τ*_*relaxM*_ and *τ*_*relaxC*_ versus grazing rate *g*. C,F: The variance *V ar*_*M*_ and *V ar*_*C*_ versus grazing rate *g*. A-C: *D* = 5.0 × 10^−4^, D-F: *D* = 1.0 × 10^−3^. (*h* = 0.44)

Our landscape-flux framework offers substantial advantages over traditional CSD-based indicators, particularly in its ability to provide earlier detection of approaching transitions. While CSD focuses primarily on local stability properties near equilibrium states, our method captures global stability characteristics and non-equilibrium dynamics across the entire state space.

Our study demonstrates that the landscape-flux approach and its derived early-warning signals (cross-correlation function Δ*CC* multidimensional data) can detect approaching transitions earlier than critical slowing down indicators based on theoretical relaxation time *τ*_*relax*_ (one-dimensional data, measured through autocorrelation). This earlier detection capability is crucial for ecological management, as it potentially provides a longer window for intervention before critical transitions occur. This comparison is particularly meaningful because relaxation time represents the fundamental dynamical property underlying all CSD indicators, rather than just comparing with empirical manifestations of CSD (such as variance or autocorrelation methods). By demonstrating advantages at this fundamental level, we establish the theoretical superiority of our approach.

Real-time ecological monitoring data from coral reef ecosystems presents an unprecedented opportunity to bridge theoretical frameworks with empirical validation. By integrating time-series data from reef monitoring stations-capturing coral cover, algal abundance, and environmental parameters-into our landscape-flux methodology, we can operationalize the theoretical results outlined above. In particular, the cross-correlation functions of the coral-reef ecosystem can be estimated directly from observed time series and hence we may calculate the average difference between forward and backward cross-correlation as an empirical EWS. Our framework thus provides practical early warning tools for policymakers and researchers, bridging the gap between abstract mathematical models and urgent conservation needs in threatened ecosystems.

## 4 Conclusions

We explored the global dynamics of a coral-algal model under stochastic fluctuations using landscape-flux theory from nonequilibrium statistical physics. In this framework, system dynamics are governed by two fundamental components: potential landscapes that guide the system toward local minima, and curl fluxes that drive transitions between alternative stable states. Quantifying global stability in complex ecological systems requires identifying an appropriate Lyapunov function-a challenging task that our approach addresses through the intrinsic potential landscape *ϕ*_0_, which serves as a Lyapunov function in the small noise limit and effectively quantifies the global stability of coral-algal dynamics.

The presence of non-zero fluxes creates a notable deviation from classical equilibrium dynamics-dominant transition paths between alternative stable states do not follow simple steepest descent trajectories on the population-potential landscape. Instead, transitions from *Macroalgae* to *Coral* states and vice versa follow irreversible paths determined by the interplay between the underlying population-potential landscape and non-zero curl fluxes. This directional path asymmetry represents a fundamental characteristic of non-equilibrium systems.

Within the bistable regime, the basin of attraction for the current state remains non-flat until reaching the right bifurcation point. Under sufficiently small noise conditions, this property enables prediction of impending state transitions before the system reaches critical points. Small fluctuations remain insufficient to trigger state switching until the right bifurcation point, where the current state’s basin flattens completely as the alternative basin becomes dominant. Consequently, time irreversibility measured through cross-correlation differences between forward and backward trajectories provides an effective predictor for approaching bifurcations, even when the system remains within its current basin without transitioning.

The analysis identifies several quantitative markers for system stability and dynamics: barrier heights between stable states, kinetic switching times (mean first passage time *MFPT*), thermodynamic cost (entropy production rate *EPR*), and dynamical driving force (average flux). We observed consistent trends across multiple metrics: average flux *Flux*_*av*_, entropy production rate *EPR*, intrinsic average flux *inFlux*_*av*_, and intrinsic entropy production rate *inEPR*. The rotational nature of flux tends to destabilize point attractors, providing a dynamical mechanism underlying phase transitions in coral-algal ecosystems. Maintaining non-equilibrium flux requires energy dissipation, revealing the thermodynamic origin of bifurcations. Intrinsic free energy also serves as an effective early warning indicator, with all these metrics exhibiting significant changes and characteristic peaks between the two transcritical bifurcations-patterns that become even more pronounced in the zero-fluctuation limit of intrinsic potential landscapes.

The nonequilibrium indicators average flux *Flux*_*av*_, entropy production rate *EPR*, time irreversibility Δ*CC*, and nonequilibrium free energy-all function as reliable predictors for critical transitions. Their turning points (peaks or troughs) consistently appear between the two transcritical bifurcations, enabling prediction of both bifurcations before the current state’s landscape flattens. These nonequilibrium warning signals precede the right bifurcation when starting from the *Macroalgae* state with increasing grazing rate *g*, and similarly anticipate the left bifurcation when starting from the *Coral* state with decreasing *g*. This bidirectional predictive capacity provides substantially earlier warnings than conventional critical slowing down theory for both bifurcation types. While specific tipping point locations may vary across different models(Xu et al., 2021, 2023), we propose that nonequilibrium indicator turning points occurring between transcritical bifurcations represent a generic feature of systems with similar qualitative dynamics.

In the current model, we utilize uncorrelated white noise as a mathematical simplification that provides analytical tractability while still capturing the essential stochastic nature of state transitions. This approach allows us to derive expressions for potential landscapes and flux patterns. We recognize that environmental disturbances like temperature fluctuations, storm events, or nutrient pulses would indeed affect coral, algae, and algal turfs in coordinated ways, introducing correlations in the noise structure of natural reef systems(Jouffray et al., 2015; Norström et al., 2009; Diko, 2010; Gardner et al., 2003; Mcmanus and Polsenberg, 2004). It is worth noting that our potential landscape-flux framework remains theoretically appropriate for systems with correlated noise. The mathematical formalism can accommodate various noise structures, including anisotropic and correlated fluctuations. Due to space limitations in the present manuscript and its complexity, we have focused on the uncorrelated case as a first approximation. The extension to correlated noise models, which would more accurately reflect synchronized environmental forcing experienced by different reef components, will be addressed in future research.

Without conducting significant further analysis, it is challenging to accurately predict the precise effects of correlated noise on the overall system dynamics. The specific correlation patterns, time scales, and amplitudes of the noise would significantly influence the system’s response. The introduction of correlation structures in stochastic perturbations fundamentally alters the statistical properties of system trajectories, potentially creating emergent behaviors that cannot be intuited through qualitative reasoning alone. The precise correlation structure to be introduced would need to be motivated by data and may differ by reef location and climate, making this a nontrivial extension of the current work but undoubtedly a valuable and interesting one.

The model tracks the evolution of proportions of space occupied by each functional type, effectively assuming that the system is spatially well-mixed, leading to a spatially implicit modeling framework (Mumby et al., 2007). This approach is appropriate for intermediate spatial scales where mixing processes (such as larval dispersal, water circulation, and mobile herbivore grazing) tend to homogenize local variations. The spatially implicit framework allows us to focus on ecosystem-level dynamics without the computational complexity of spatially resolved models. Our potential and flux field landscape theoretical framework offers considerable versatility and could be naturally extended to spatially explicit models in future research. We recognize the importance of spatial heterogeneity in coral reef ecosystems and in subsequent work, we plan to develop spatially explicit extensions of this framework. In recent work, we have shown that the framework can be extended to spatially explicit models (of vegetation dynamics) and hence it is a natural next step to leverage this progress to explore how the present results compare with EWSs in spatial extension of the coral reef model studied here(Siu et al., 2025; Wu and Wang, 2013b, 2014, 2013a; Lepzelter and Wang, 2008).

Despite the simplifying assumptions of the mathematical model, our current framework provides valuable insights into the global stability of coral reef ecosystems and demonstrates the utility of landscape-flux theory for understanding complex ecological dynamics. The simplifications employed here serve as a necessary first step toward more comprehensive models that can incorporate the full complexity of coral reef systems (Mcmanus et al., 2018; Nes et al., 2016).

The ongoing degradation of coral reefs and deterioration of reef ecosystems remain among the most pressing conservation challenges of our time. By advancing theoretical understanding of coral-algae dynamics through our potential landscape-flux approach, this study contributes valuable insights that may guide practical conservation strategies for protecting and restoring these ecologically crucial yet increasingly threatened marine ecosystems.

### Data availability

All study data are included in this article and/or the supplemental information. Any additional information required is available from the corresponding authors contact upon request.

## Appendix A

Potential landscape and Local Stability Analysis of Equilibrium Systems

In equilibrium systems, the potential function or landscape is an essential tool for describing the stability of system states. For such systems, dynamics are completely determined by the potential landscape, with the system always evolving along the direction of decreasing potential energy until reaching a potential energy minimum. A key characteristic of equilibrium systems is the absence of non-zero probability flux, meaning the system satisfies detailed balance conditions, with zero net flow along any closed path being zero (Wang, 2015; Ge and Qian, 2010; Qian, 2006; Nicolis and Prigogine, 1977; Van Kampen, 2007; M., 1992).

Mathematically, the dynamic equation of an equilibrium system can be represented as a gradient system: *dx/dt* = −∇*U* (*x*), where *U* (*x*) is the potential function landscape. The system’s steady states correspond to extremal points of the potential function, with minima representing stable equilibrium points and maxima representing unstable equilibrium points.

Local stability analysis is a method for studying the behavior of small perturbations near equilibrium points. By linearizing the dynamic equations around an equilibrium point, one obtains the Jacobian matrix characterizing the fluctuations. For equilibrium systems, this matrix is symmetric, and its eigenvalues completely determine the stability of the equilibrium point (Nicolis and Prigogine, 1977; Van Kampen, 2007; M., 1992): - All negative eigenvalues: stable node - Presence of positive eigenvalues: unstable equilibrium point - Presence of zero eigenvalues: potential bifurcation

The potential landscape of equilibrium systems visually demonstrates the global stability structure of the system, with low potential energy regions corresponding to states where the system is more likely to reside, while the height of potential barriers reflects the difficulty of state transitions. This analytical approach has wide applications in the study of physical, chemical, and biological systems.

## Author contributions

SL and JW designed the conceptualization and LX and DP carried them out. LX, DP, SL and JW developed the model code and performed the simulations. LX, DP, SL and JW prepared the manuscript.

## Acknowledgements

LX thanks supports by Natural Science Foundation of Jilin Province No. 20220101013JC and National Natural Science Foundation of China No.12234019. SL and DP appreciate support from NSF DMS-1951358.

