## Supplemental Table 1 for "Global Stability and Tipping Point Prediction in a Coral-Algae Model Using Landscape-Flux Theory"

#### Non-equilibrium thermodynamics, entropy, energy and free energy of the general dynamical systems under the zero-fluctuation limit and the finite fluctuations

The intrinsic potential $\phi_0$ and the steady-state probability distribution in non-equilibrium systems have the relationship as follows: $\mathcal{P}_{ss}(\mathbf{x}) = P_{ss}(\mathbf{x})|_{D\to 0} = \exp(-\phi_0/\mathcal{D})/\mathcal{Z}$, where $\mathcal{D} = D|_{D\to 0}$. The partition function is $\mathcal{Z} = \int \exp(-\phi_0/\mathcal{D}) d\mathbf{x}$. $\phi_0$ is $\phi_0 = -\mathcal{D}\ln(\mathcal{Z}\mathcal{P}_{ss})$.

$\mathcal{S}$ is the entropy of the non-equilibrium system under the zero-fluctuation limit as $\mathcal{S} = -\int \mathcal{P}(\mathbf{x},t)\ln\mathcal{P}(\mathbf{x},t)d\mathbf{x}$ (Wang, 2015; Zhang et al., 2012; Xu et al., 2014; Wang et al., 2008; Zhang et al., 2012).

where $\mathcal{E} = \int \phi_0 \mathcal{P}(\mathbf{x},t)d\mathbf{x} = -\mathcal{D}\int \ln(\mathcal{Z}\mathcal{P}_{ss})\mathcal{P}(\mathbf{x},t)d\mathbf{x}$ is the intrinsic energy of the non-equilibrium system. And $\mathcal{F} = \mathcal{E} - \mathcal{D}\mathcal{S} = \mathcal{D}\left(\int \mathcal{P}\ln(\mathcal{P}/\mathcal{P}_{ss})d\mathbf{x} - \ln\mathcal{Z}\right)$ is the intrinsic free energy of the non-equilibrium system.

We can calculate the time evolution of the intrinsic free energy as $\frac{d\mathcal{F}}{dt} = -\mathcal{D}^2\left(\int \left[\nabla\ln(\frac{\mathcal{P}}{\mathcal{P}_{ss}})\cdot\mathbf{G}\cdot\nabla\ln(\frac{\mathcal{P}}{\mathcal{P}_{ss}})\right]\mathcal{P}d\mathbf{x}\right) \leq 0$ (Zhang et al., 2012; Xu et al., 2014). This implies that $\mathcal{F}$ always decreases and it is a Lyapunov function. It can be used to quantify the global stability of the non-equilibrium system at finite fluctuations. $\mathcal{F}$ has its minimum with $\mathcal{F} = -\mathcal{D}\ln\mathcal{Z}$.

#### Kinetic speed and dominant paths between the $Macroalgae$ state and the $Coral$ state

We can also use the path-integral approach to identify the dominant paths between stable states. The probability of the path from initial state $\mathbf{x}_i$ at $t=0$ to final state $\mathbf{x}_f$ at time $t$ follows is given below with the Lagrangian $L(\mathbf{x}(t))$ and the action $A(\mathbf{x})$ (Wang et al., 2011; Xu et al., 2014): $P(\mathbf{x}_f,t|\mathbf{x}_i,0) = \int D\mathbf{x}\ \exp[-\int dt(\frac{1}{2}\nabla\cdot\mathbf{F}(\mathbf{x}) + \frac{1}{4}(d\mathbf{x}/dt - \mathbf{F}(\mathbf{x}))\cdot(D\mathbf{G})^{-1}\cdot(d\mathbf{x}/dt - \mathbf{F}(\mathbf{x})))] = \int D\mathbf{x}\exp[-A(\mathbf{x})] = \int D\mathbf{x}\exp[-\int L(\mathbf{x}(t))dt]$.

$D\mathbf{x}$ denotes the sum over all possible paths connecting $\mathbf{x}_i$ at time zero to $\mathbf{x}_f$ at time $t$(Wang et al., 2011; Xu et al., 2014). We can obtain the dominant paths with the optimal weights by minimization of the weight or the action $A(\mathbf{x})$.

### A visual analogies for mathematical concepts

Our approach reveals coral-algae dynamics through two complementary concepts: landscape and flux.

The Landscape can be visualized as a physical terrain with hills and valleys. Like a marble rolling on this surface, coral-algae systems will naturally move toward valleys (stable states) and away from hills (unstable states). The depth of valleys indicates stability strength-deeper valleys represent more resilient states that withstand stronger disturbances. Conversely, the height of the barrier between valleys determines transition difficulty.

The Flux is analogous to river currents creating circular patterns that can't be explained by downhill movement alone. Just as water doesn't flow directly downhill but forms eddies and swirls, flux creates rotational movements in the system dynamics; in contrast to the influence of the landscape, which pushes the system directly towards stable states. The flux is also responsible for time irreversibility, a fundamental characteristic of complex systems. In terms of the river metaphor, if you drop a leaf in the water and watch its journey downstream, you'll observe a specific path, but trying to reverse this journey by pushing the leaf back upstream won't mirror the original path exactly. This irreversibility occurs because eddies and currents create asymmetric flow patterns, the leaf encounters different obstacles in different sequences during forward versus backward movement. Similarly, the flux is responsible for the fact that the most likely coral-to-algae transitions follow different paths than algae-to-coral recoveries in our model. These circular patterns require continuous energy input to maintain-another key characteristic of non-equilibrium systems.

Together, these complementary forces of landscape and flux provide a complete picture of coral-algae behavior. The landscape determines where stable states exist and their resistance to perturbations, while flux shapes the actual transition pathways between alternative stable states, creating the rich, sometimes surprising dynamics observed in these complex systems.

Entropy production measures how costly it is for a system to maintain its non-equilibrium state-essentially its "operating cost." As a system approaches a critical transition, this "cost" often changes significantly. By measuring entropy production, we can detect when a coral-algae system is working "harder" to maintain its current state, potentially signaling an upcoming transition even before visible changes occur. The $Flux_{av}$ represents the average strength of the circular flow patterns in the coral-algae system. Like measuring the average current strength in a river, $Flux_{av}$ quantifies the intensity of these non-equilibrium dynamics.

### The differences between global stability and local stability

The difference between global stability and local stability relates to the scope of a system's response to perturbations.

Local stability refers to how a system behaves when subjected to small perturbations around a specific equilibrium point. A locally stable equilibrium will return to its original state when slightly disturbed. This analysis typically involves linearization techniques and examining eigenvalues of the Jacobian matrix at the equilibrium point.

Global stability concerns the system's behavior under perturbations of any magnitude, regardless of how far the system is displaced from equilibrium. A globally stable equilibrium will eventually attract all trajectories from anywhere in the state space.

Local stability analysis provides less comprehensive information about the system's overall behavior compared to global stability analysis. Local stability only tells us about behavior in a small neighborhood around an equilibrium, while global stability provides stronger guarantees about the system's behavior across its entire state space. In dynamical systems like ecological models, local stability is often easier to determine mathematically (through eigenvalue analysis), while global stability typically requires more sophisticated techniques such as Lyapunov functions or extensive numerical simulations.

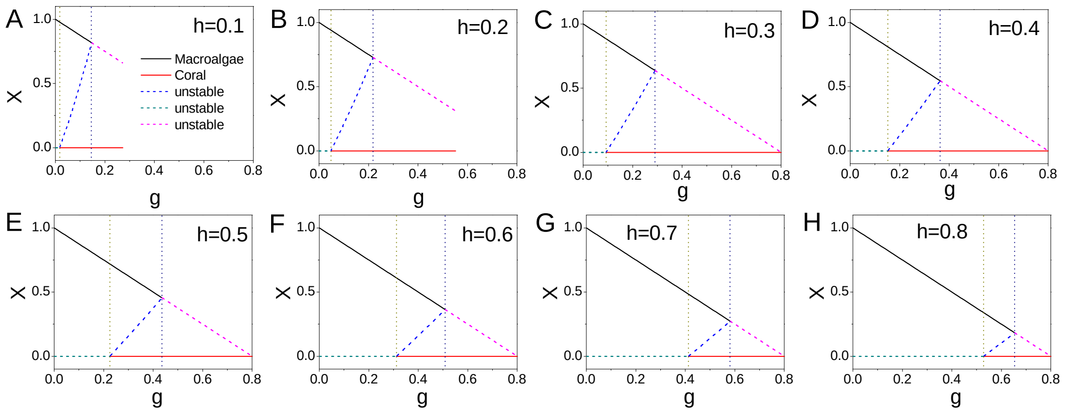


**Figure 1.** The phase diagram versus grazing rate $g$ with different natural mortality rate of corals $h$ for the algae-coral model.

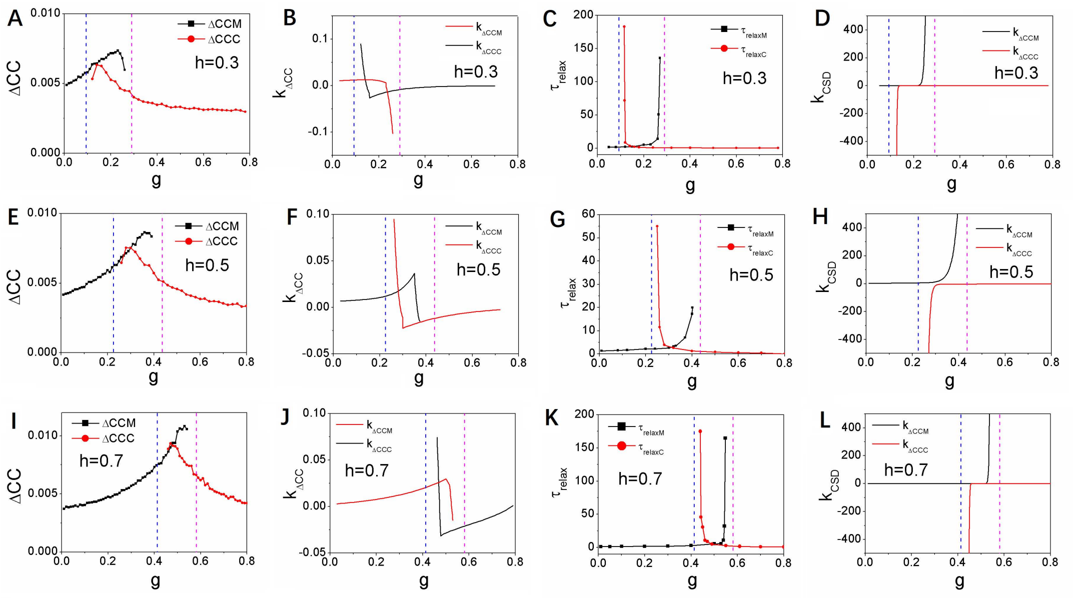


**Figure 2.** A(E,I): The average difference of the cross correlations forward and backward in time $\Delta CCM$ (for state $Macroalgae$) and $\Delta CCC$ (for state $Coral$)versus $g$. B (F,J): $k_{\Delta CCM}$ (the slope of $\Delta CCM$) and $k_{\Delta CCC}$ (the slope of $\Delta CCC$) versus $g$. C (G,K): The relaxation time $\tau_{relaxM}$ and $\tau_{relaxC}$ versus grazing rate $g$. D (H,L): $k_{CSDM}$ (the slope of the relaxation time $\tau_{relaxM}$) and $k_{CSDC}$ (the slope of the relaxation time $\tau_{relaxC}$) versus grazing rate $g$. A-D:$h = 0.3$, E-H:$h = 0.5$, I-L:$h = 0.7$.

| Term | Definition |
| --- | --- |
| Attractor State | A stable configuration toward which a dynamical system tends to evolve regardless of starting conditions within its basin of attraction. |
| Basin of Attraction | The set of initial conditions from which a dynamical system evolves toward a particular attractor state. |
| Barrier height ($\Delta U$,$\Delta \phi_0$) | energy required for a population to transit from one stable state to another in ecological or biological systems. |
| Bifurcation | A qualitative change in system behavior that occurs when a small change in a parameter causes a sudden structural change in the system's dynamics. |
| Bistable Region | A parameter range where two stable states (attractors) coexist, allowing the system to settle in either state depending on initial conditions and perturbations. |
| Critical Slowing Down (CSD) | The phenomenon where a system takes progressively longer to recover from perturbations as it approaches a bifurcation point, manifested as increasing relaxation time. |
| Cross-Correlation Function | A statistical measure of similarity between two time series as a function of the displacement of one relative to the other, used to quantify temporal relationships. |
| Curl Flux ($J$) | The rotational component of system dynamics that breaks detailed balance and drives non-equilibrium behavior, causing circulation in state space. |
| Detailed Balance | A condition in equilibrium systems where the probability flux between any two states is balanced, resulting in zero net circulation. |
| Diffusion Coefficient ($D$) | A parameter quantifying the intensity of random fluctuations (noise) in a stochastic system. |
| Dominant Path | The most probable trajectory a system follows when transitioning between stable states. |
| Entropy Production Rate ($EPR$) | A thermodynamic quantity measuring the rate at which a non-equilibrium system dissipates energy and generates entropy, quantifying the thermodynamic cost of maintaining non-equilibrium dynamics. |
| Frequency of the flickering ($f_\omega$) | the number of transitions from one stable state to another stable state per unit time. |
| Inflection Point | A point on a curve where the curvature changes sign, used to identify significant changes in system behavior. |
| Intrinsic Free Energy ($\mathcal{F}$) | The free energy of a system calculated in the zero-noise limit, representing fundamental system energetics. |
| Intrinsic Potential Landscape ($\phi_0$) | The potential landscape in the zero-noise limit that serves as a Lyapunov function for determining global stability. |
| Landscape-Flux Theory | A theoretical framework for analyzing non-equilibrium systems by decomposing dynamics into gradient (landscape) and rotational (flux) components. |
| Langevin Equation | A stochastic differential equation describing the time evolution of a system subject to random forces. |
| Lyapunov Function | A scalar function that quantifies a system's stability, decreasing along trajectories and reaching minima at stable equilibria. |
| Mean First Passage Time ($MFPT$,$\tau$) | The average time required for a system to transition from one stable state to another, quantifying kinetic switching times between states. |
| Natural Mortality Rate of Coral ($h$) | A parameter representing the intrinsic death rate of coral in the coral-algal model. |
| Non-Equilibrium System | A system that does not satisfy detailed balance conditions and requires continuous energy input to maintain its state. |
| Phase Diagram | A graphical representation showing different stable states and transition boundaries as functions of control parameters. |
| Potential Landscape ($U$) | A mathematical function whose gradient determines the force driving a system toward stable states, analogous to an energy surface. |
| Relaxation Time ($\tau_{relax}$) | The characteristic time required for a system to return to equilibrium after a perturbation. |
| Saddle Node Bifurcation | A bifurcation where a stable and an unstable fixed point collide and annihilate each other. |
| Tipping Point | A critical threshold where a small change in a parameter causes a rapid, often irreversible transition to a qualitatively different system state. |
| Time Irreversibility ($\Delta CC$) | A measure quantifying asymmetry between forward and backward processes in time, indicating non-equilibrium behavior and serving as an early warning signal. |
| Transcritical Bifurcation | A bifurcation where two fixed points (one stable, one unstable) exchange stability as a parameter changes. |

**Table 1.** Glossary of terms used in landscape-flux theory
